# CDK4/6 inhibition enhances SHP2 inhibitor efficacy and is dependent upon restoration of RB function in malignant peripheral nerve sheath tumors

**DOI:** 10.1101/2023.02.02.526674

**Authors:** Jiawan Wang, Ana Calizo, Lindy Zhang, James C. Pino, Yang Lyu, Kai Pollard, Xiaochun Zhang, Alex T. Larsson, Eric Conniff, Nicolas Llosa, David K. Wood, David A. Largaespada, Susan E. Moody, Sara J. Gosline, Angela C. Hirbe, Christine A. Pratilas

## Abstract

Malignant peripheral nerve sheath tumors (MPNST) are highly aggressive soft tissue sarcomas with limited treatment options, and novel effective therapeutic strategies are desperately needed. We observe anti-proliferative efficacy of genetic depletion or pharmacological inhibition using the clinically available SHP2 inhibitor (SHP2i) TNO155. Our studies into the signaling response to SHP2i reveal that resistance to TNO155 is partially mediated by reduced RB function, and we therefore test the addition of a CDK4/6 inhibitor (CDK4/6i) to enhance RB activity and improve TNO155 efficacy. In combination, TNO155 attenuates the adaptive response to CDK4/6i, potentiates its anti-proliferative effects, and converges on enhancement of RB activity, with greater suppression of cell cycle and inhibitor-of-apoptosis proteins, leading to deeper and more durable anti-tumor activity in *in vitro* and *in vivo* patient-derived models of MPNST, relative to either single agent. Overall, our study provides timely evidence to support the clinical advancement of this combination strategy in patients with MPNST and other tumors driven by loss of NF1.

## INTRODUCTION

Malignant peripheral nerve sheath tumors (MPNST) are highly aggressive heterogeneous soft tissue sarcomas with limited effective treatment options, given their relative insensitivity to conventional systemic chemotherapy and radiation, and their propensity to metastasize. In many patients with neurofibromatosis type 1 (NF1), MPNST develop from within plexiform neurofibromas (pNF) (*1*), the benign precursor tumors which themselves can be sources of pain, disfigurement, and alteration of function (*2, 3*). The most recurrent genomic alterations underlying the pathogenesis of MPNST are loss of function (LOF) of the RAS-GAP (GTPase-activating protein) neurofibromin [NF1, 90%, ref. (*4, 5*)], *CDKN2A* (60-80%), PRC2 [70-90%, ref. (*4–7*)], and *TP53*, as well as gain of chromosome 8 [80%, ref. (*8, 9*)] yet, molecular targeting of these LOF alterations represents a unique challenge and a major unmet need for these patients.

Despite many clinical trials of chemotherapy and targeted agents, there has been little advancement in overall survival for patients with MPNST, and thus novel therapeutic approaches are desperately needed (*10*). Currently, only one class of drug, MEK inhibitors (MEKi), has been approved for reducing symptomatic tumor burden in patients with pNF (*11–13*). Aberrantly upregulated ERK signaling, due to loss of NF1 and dysregulated PRC2 function, is a critical effector of tumorigenesis, and thus pharmacological MEK inhibition has also been tested in models of MPNST (*11, 14–16*). However, the preclinical responses of MPNST to monotherapy with MEKi and other targeted agents are incomplete, suggesting a need for improved understanding of the role of ERK and other RAS effector pathways (*17, 18*).

The incomplete effects of MEKi in MPNST prompted us to explore potential combinatorial therapeutics using MEKi and agents targeting the adaptively changed signaling elements that emerge upon short-term MEK inhibition. We have previously reported that the adaptive and acquired response to MEKi in MPNST involves the upregulation of activity of multiple receptor tyrosine kinases (RTK) (*19, 20*). SHP2 represents a promising target and inhibition of SHP2 can target a point of convergence from upstream RTK signaling, and together with inhibition of RAS effector pathways provide an enhanced anti-tumor effect (*11, 21–24*). SHP2, encoded by *PTPN11*, facilitates RAS-GEF-mediated RAS-GTP loading and subsequent membrane recruitment of RAS and therefore is required for RTK-mediated RAS/ERK activation (*23*). SHP2 phosphatase also acts as a positive regulator of RAS by dephosphorylating RAS and promoting its binding to and activation of RAF kinases, leading to increased ERK signaling (*25*). Thus, we tested combination SHP2i as a strategy to overcome RTK upregulation seen with single agent MEKi in tumors with loss of NF1, and subsequently demonstrated that combined MEKi/ SHP2i is additive in both *in vitro* and *in vivo* models of NF1-MPNST (*20*).

We then observed more potent single agent activity of the clinically available SHP2i TNO155 (NCT04000529), compared to SHP099, and therefore hypothesized that SHP2i, rather than MEKi, may be a critical anti-tumor strategy for NF1-MPNST. As potential clinical therapies for MPNST continue to evolve in response to both clinical and preclinical data, MEKi may not in fact be the most ideal combination partner for two reasons: 1) many patients who develop MPNST will have already been exposed to MEKi as treatment for their precursor pNF, and 2) SHP2i and MEKi may result in overlapping toxicity profiles due to ERK signaling suppression as an effect of both agents, leading to poor tolerability in people. In fact, in the ongoing study of a SHP2i and a MEKi (NCT03989115), drugs are administered on an intermittent dosing schedule to address toxicity (*26*).

We therefore sought an alternate partner to combine with SHP2i in order to advance this agent to a clinical trial for people with MPNST, and we have focused specifically on CDK4/6i. Frequent loss of *CDKN2A*, inactivation of RB, hyperactivation of cyclin dependent kinases (CDK), and the dependency of D-cyclins on RAS signaling suggest that CDK4/6i may be an attractive therapeutic strategy in NF1-MPNST (*27, 28*). We hypothesized that the cytostatic effects of CDK4/6i may be potentiated by drugs targeting upstream regulators of RAS, such as SHP2, in MPNST.

In this study, we demonstrate that TNO155 plus the CDK4/6i ribociclib produces additive anti-proliferative and anti-tumor effects in cell lines and patient-derived xenograft (PDX) models of NF1-MPNST through synergistic inhibition of cell cycle and induction of apoptosis. When given together, TNO155 attenuates the adaptive response to CDK4/6i, produces deeper and more durable responses, and thus potentiates the efficacy of single agent CDK4/6i in a broad collection of preclinical models that are a faithful representation of the genomically heterogenous human MPNST. Taken together, with the clinical testing of TNO155 and ribociclib currently underway (NCT04000529), this study will serve as an important contribution to the advancement of novel SHP2i combinations in the clinic for people with NF1-deficient cancers.

## RESULTS

### *PTPN11* genetic depletion reduces MPNST cell growth and alters MEK/ERK activity

Activating mutations in *PTPN11*, the gene that encodes SHP2, occur in about 50% of patients with Noonan syndrome and a subset of leukemias (*29*), whereas in solid tumors, *PTPN11* mutations occur at a low frequency. Wild-type (WT) SHP2 is critical for activation of RTK/RAS/ERK signaling which drives proliferation and growth of tumor cells. To understand the functional role of WT SHP2 in NF1-MPNST growth, we knocked down SHP2 by generating doxycycline-inducible expression of two unique shRNA targeting *PTPN11* (#818, and #5003) in three NF1-MPNST cell lines (Fig 1 and Fig S1) including the two traditional lines ST8814, NF90.8 and one patient derived MPNST cell line JH-2-002 (*30*). *PTPN11*/ SHP2 genetic depletion significantly reduced MPNST cell growth, as evidenced by long-term colony formation assay (Fig 1A and Fig S1A) and short-term real-time cell confluence monitoring, using Incucyte imaging system (Fig 1B and Fig S1B). SHP2 protein expression and phosphorylation at tyrosine 542 were dramatically decreased following SHP2 knockdown after 72-hour treatment with doxycycline (Dox), while the signaling intermediates phospho-MEK (p-MEK) and phospho-ERK (p-ERK) exhibited marginal reduction (Fig 1C and Fig S1C), likely due to rebound in ERK signaling.

**Fig. 1.**
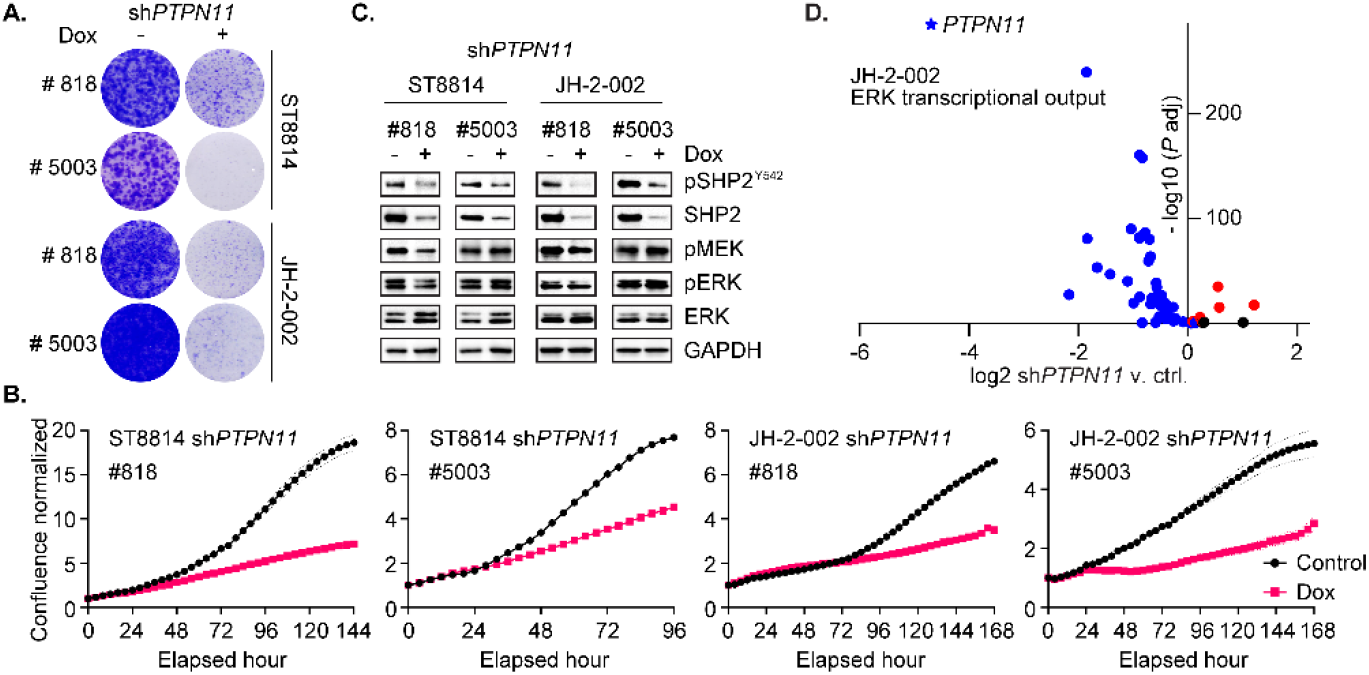
*PTPN11* genetic depletion reduces MPNST cell growth and alters MEK/ERK activity. (**A**) ST8814 and JH-2-002 transduced with doxycycline (Dox) inducible constructs expressing sh*PTPN11* #818 or #5003, were treated with vehicle or 300 ng/ml Dox for two weeks. Cells were fixed with 10% neutral buffered formalin and then stained with crystal violet. (**B**) Cells as in A. were treated with vehicle or 300 ng/ml Dox for up to seven days. The phase confluence was monitored by Incucyte real-time imaging system, normalized to corresponding 0-hour scan. (**C**) Cells as in A. were treated with vehicle or 300 ng/ml Dox for 72 hours, and the indicated proteins were assessed using immunoblot. (**D**) JH-2-002 transduced with Dox-inducible sh*PTPN11* #818 was treated with vehicle or 300 ng/ml Dox for 72 hours, and then three replicates were collected for RNAseq, and the fourth replicate was collected for immunoblotting validation. RNAseq result revealed a significant decrease in *PTPN11* RNA expression (n= 3, *P* adj= 0, LFC= −4.69) when comparing sh*PTPN11* v. ctrl. A total of 51 genes representing MEK/ERK transcriptional output (*31*), were assessed for their expression after *PTPN11* knockdown. A volcano plot demonstrating log2 fold change of sh*PTPN11* v. ctrl as a function of −log10 (*P* adj) is shown. Black dots= not significant (ns, *P* adj> 0.05); red dots= log2 fold change (LFC)> 0 and blue dots= LFC< 0 (*P* adj< 0.05). 73% (37/51) of genes in the set were significantly transcriptionally downregulated following *PTPN11* genetic depletion.

We have previously found that the level of p-ERK, which has been used as a surrogate marker for RAS/ERK pathway activation, is inherently susceptible to feedback regulation by DUSPs, and therefore not an accurate measure of signaling output (*31*). MEK/ERK-dependent transcriptional output is defined as the set of genes whose expression changes significantly after 8-hour MEK inhibition (*31*). In JH-2-002 sh*PTPN11*#818, treated with Dox or vehicle, we first validated protein expression level of SHP2 and other signaling proteins (Fig S1D), and performed RNA sequencing (RNAseq, see Materials and Methods). We confirmed significant reduction in transcript levels of *PTPN11* and analyzed the transcriptional profile of sh*PTPN11* relative to defined ERK signaling output (Fig 1D). A total of 51 genes (excluding *KIR3DL2*, which was not detected via RNAseq), are included in the analysis (Table S1). 73% (37/51) of genes in the MEK-dependent set demonstrated significant downregulation after *PTPN11* depletion (*P* adj <0.05), indicating a global response of ERK signaling suppression by SHP2 knockdown (Fig 1D and Fig S1E). Taken together, these observations concluded that *PTPN11* genetic depletion reduces MPNST cell growth in an ERK signaling dependent manner.

### SHP2i TNO155 alters growth and gene expression in NF1-MPNST cells

We first characterized the genomic or protein alterations in *NF1*, *CDKN2A*, PRC2 components *SUZ12* and *EED*, *TP53*, and others of importance in an updated panel of MPNST cell lines [Fig 2A, ref. (*19*)] to confirm that our cell-line panel is a faithful genomic representation of the highly heterogeneous MPNST seen in patients. We next tested the effect of pharmacological inhibition of SHP2 on NF1-MPNST cell proliferation, viability, and growth. While SHP099 (a tool compound which has not been tested in clinical trials) was only partially effective in reducing cell and tumor growth in our initial studies (*20*), we found that the lead clinical compound, TNO155, which is now being tested in clinical trials (NCT03114319, NCT04000529), was more potently able to suppress cell proliferation and growth in 10 MPNST *in vitro* models (Fig 2B and C). SHP099 also demonstrated activity against cellular viability in a 3D microtissue *ex-vivo* model derived from four unique MPNST PDX [Fig S2A, ref. (*9*)]. Further investigation in ST8814 and JH-2-079c cell lines (*9*) treated with DMSO or 0.3 μmol/L TNO155, using Incucyte real-time monitoring system, demonstrated caspase-3/7 activation induced by TNO155 (Fig S2B). Mechanistically, we found that TNO155 inhibited SHP2 phosphatase activity as evidenced by p-SHP2 reduction, and markedly suppressed ERK signaling and downstream cell cycle pathways, in a dose-dependent (Fig 2D) and time-dependent (Fig 2E) manner. To further confirm the effect of TNO155 on ERK signaling and cell cycle regulators, we expanded the investigation to a collection of eight NF1-MPNST cell lines including three patient-derived lines (JH-2-002, JH-2-079c and JH-2-103). We again observed that TNO155 effectively inhibited cell cycle regulators via ERK signaling suppression (Fig 2F). Notably, TNO155 induced RB1 hypo-phosphorylation consistently across the cell lines tested, in a dose- and time-dependent manner (Fig 2D-F).

**Fig. 2.**
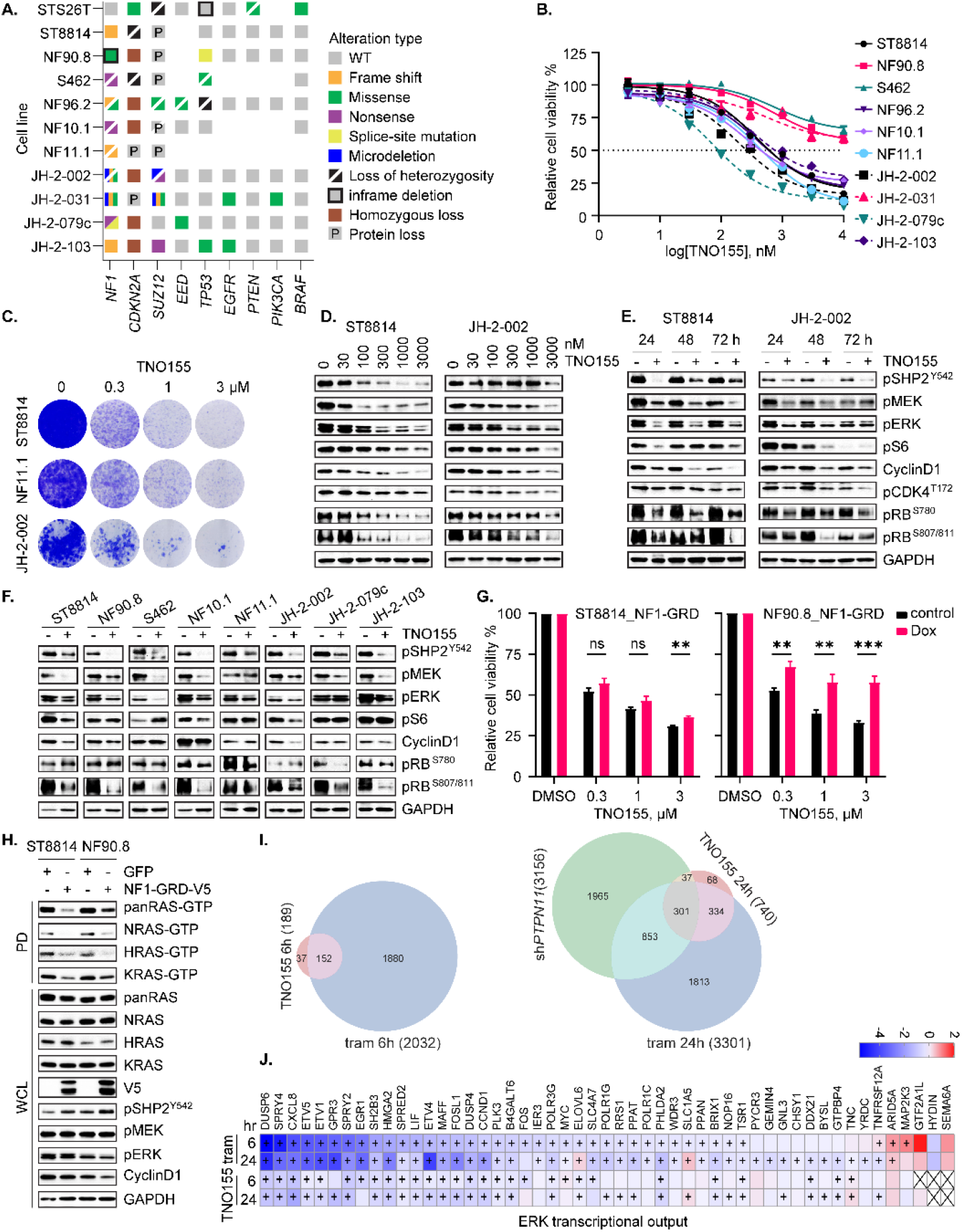
SHP2i TNO155 alters growth and gene expression in NF1-MPNST cells. (**A**) Onco-print of key driver genes (*NF1, CDKN2A, SUZ12* and *EED*) in MPNST and putative others across a panel of MPNST cell lines is shown. (**B**) Ten NF1-associated MPNST cell lines were treated with increasing dose of the SHP2i TNO155 for five days. Cell viability was evaluated by using the cell counting kit-8 (CCK-8) assay measuring metabolic activity. (**C**) Three NF1-MPNST cell lines were treated with DMSO or TNO155 (0.3, 1 and 3 μM) for about two weeks. Cells were fixed with 10% neutral buffered formalin and then stained with crystal violet. (**D**-**E**) ST8814 and JH-2-002 were treated with increasing dose (30-3000 nM) of TNO155 for 24 hours (**D**) or 0.3 μM TNO155 over a time course (**E**). Signaling intermediates in ERK and cell cycle pathways were detected using immunoblot. (**F**) Eight NF1-MPNST cell lines were treated with DMSO or 0.3 μM TNO155 for 48 hours. The representative signal intermediates in ERK and cell cycle pathways were detected using immunoblot. (**G**) ST8814 and NF90.8 transduced with doxycycline-inducible NF1-GRD (GAP related domain) were pretreated with vehicle control or 300 ng/ml Dox, followed by treatment with DMSO or TNO155 (0.3, 1 and 3 μM) for five days. Cell viability was assessed using CCK-8. (**H**) ST8814 and NF90.8 transduced with doxycycline-inducible GFP or V5 tagged NF1-GRD were treated with 300 ng/ml Dox for 24 hours. RAS activity and signaling intermediates were detected using immunoblot following the active RAS pull-down assay. PD= pull down; WCL= whole cell lysate. (**I**) Venn diagram showing the number of overlapped significant genes (*P* adj<0.05 and |fold change|>1.5) between sh*PTPN11*, TNO155 and trametinib at 6- and 24-hour time points. (**J**) Heatmap demonstrating changes in transcriptional output of the 51 ERK signature genes, derived from RNAseq analysis of JH-2-002 after DMSO, 0.3 μmol/L TNO155 or 20 nmol/L trametinib treatment (6 and 24 hours).

Considering the competitive role of NF1 GAP function and SHP2 inhibition in reducing the accumulation of RAS-GTP during nucleotide cycling, we asked whether NF1 GAP function played a role in response to TNO155. We therefore reconstituted expression of NF1-GRD (GAP-related domain) in two traditional NF1 LOF cell lines, ST8814 and NF90.8, using a Dox-inducible murine retrovirus expression system. Intriguingly, re-introduction of NF1-GRD to restore GAP function partially rescued cell viability upon TNO155 treatment (Fig 2G), suggesting that NF1 loss confers sensitivity to SHP2i, an observation consistent with previous studies (*23, 32*). Indeed, relative to the GFP control, re-expression of NF1-GRD reduced RAS-GTP levels, indicative of a successful restoration of NF1 GAP function (Fig 2H). When comparing the cell viability and ERK signaling profiles of ST8814 versus NF90.8, the extent of resistance to TNO155 conferred upon restoration of NF1 GAP function is associated with the degree of suppression of ERK signaling and downstream effectors by NF1-GRD (Fig 2G and H).

To compare the transcriptional effects of sh*PTPN11*, TNO155, and the MEKi trametinib individually, we conducted RNAseq analysis of JH-2-002 treated with DMSO, TNO155 or trametinib at 6- and 24-hour time points, and sh*PTPN11*, in order to measure both direct and adaptive responses to ERK signaling inhibition. Analysis of the transcriptional response showed that treatment with 0.3 μmol/L TNO155 for 6 and 24 hours led to fewer differentially expressed genes compared to the treatment of 20 nmol/L trametinib or sh*PTPN11* (Fig 2I and Table S2), which is in line with the immunoblots measuring ERK signaling and cell cycle regulators at the same time points (Fig S2C). Specifically, 740 genes were significantly altered (*P* adj<0.05, absolute fold change >1.5) following TNO155, while 3301 and 3156 genes were significantly altered after trametinib and sh*PTPN11*, respectively. Most of the genes (~80%) significantly altered by TNO155 overlapped with those altered by trametinib (Fig 2I). For both treatments, the number of significantly altered genes increased over a 6- to 24-hour time course, indicating the existence of indirect and adaptive changes to transcription after ERK signaling suppression. We also assessed ERK signaling using the 51 ERK signature genes described above and found most genes in the signature to be significantly down-regulated upon drug treatment. This suggests that the TNO155 treatment, while affecting fewer genes (Fig 2I), has a pronounced effect on the ERK signature genes, despite less potent inhibition by TNO155 relative to trametinib (Fig 2J and Fig S2D, Table S1), consistent with the degree of ERK signaling inhibition by both agents observed in Figure S2C.

### Loss of function in RB1 confers resistance to TNO155

The recent study highlights the transcriptional upregulation of cell cycle regulators during the progression from pNF to MPNST (*27*). We also observed a consistently enhanced cell cycle transcriptional profile in samples from our NF1 biospecimen repository, using the gene category defined as cell cycle regulators (*27*), and found that *PLK1*, *CCNA2*, and *BIRC5* demonstrated significant upregulation in MPNST compared to pNF (Fig 3A), suggesting a driver role of cell cycle dysregulation in tumor pathogenesis and disease progression. Given the reported genomic heterogeneity of NF1-MPNST and the impact this heterogeneity has at the protein level (*19*), we sought to determine whether steady-state expression level of a specific molecular marker can predict sensitivity to SHP2i treatment. We first characterized the signaling protein markers in the updated panel of MPNST cell lines (Fig 3B and Fig S3A-B). We found that of the 9 cell lines in this panel (which also includes the sporadic MPNST line STS26T), all exhibited loss of p16 expression (Fig 3B), 7 exhibited loss of SUZ12 expression (Fig S3A), and all expressed various levels of p53 protein (Fig S3A) despite known mutations in some lines (Fig 2A). The two NF1-MPNST lines that retained SUZ12 expression (Fig S3A), NF96.2 and JH-2-079c, harbor p.S241R and p.D243Y mutations in *EED*, respectively (Fig 2A). We examined the protein profiles involved in cell cycle regulation (Fig 3B and Fig S3A) and in AKT and ERK pathways (Fig S3B) to determine if there were some alterations in baseline expression of a single unique molecule that explained the lower TNO155 sensitivity exhibited in three of the cell lines in Figure 2B, but there were no salient correlations.

**Fig. 3.**
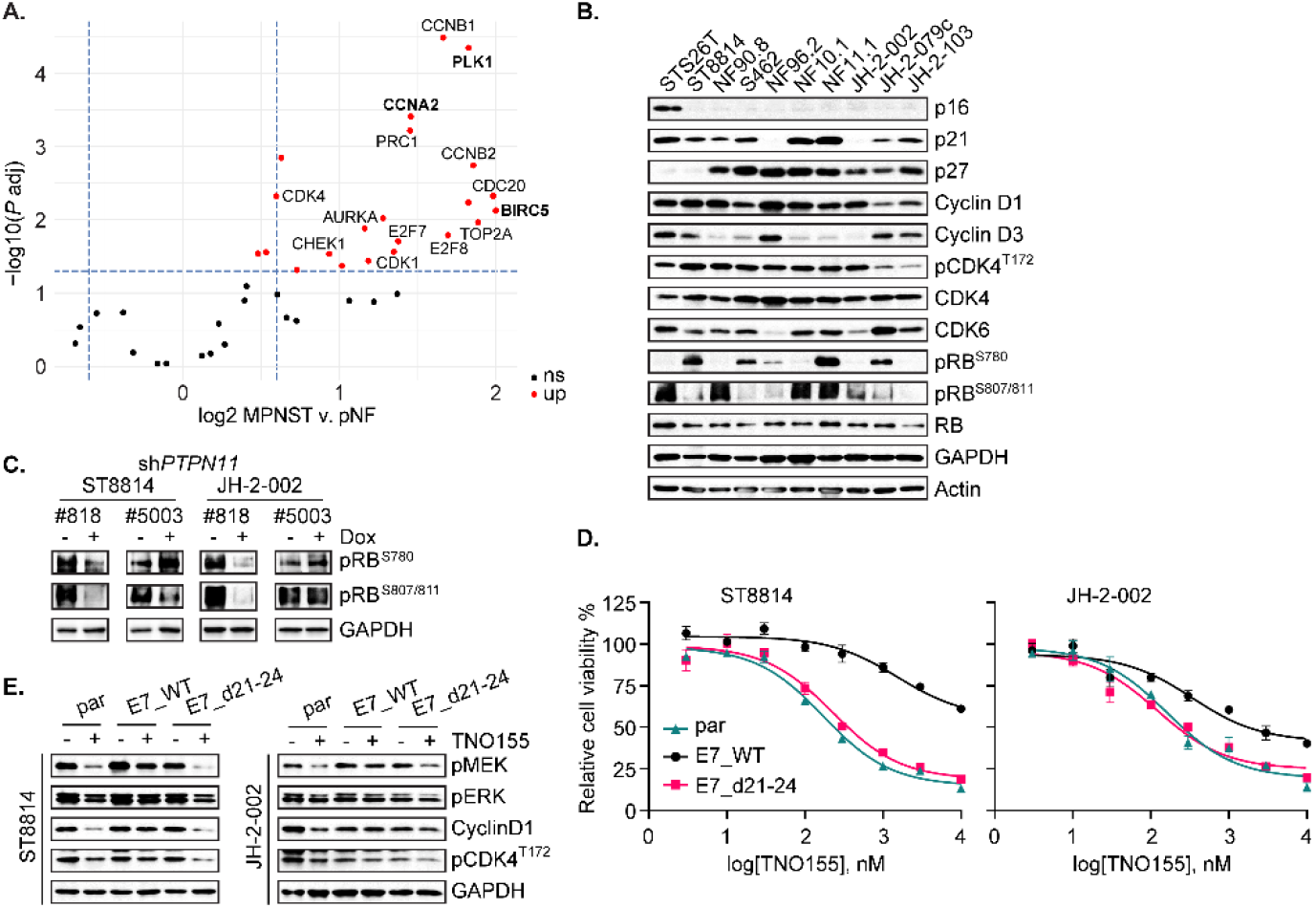
Loss of function in RB1 confers resistance to TNO155. (**A**) Volcano plot from RNAseq analysis revealing cell cycle dysregulation comparing MPNST v. pNF. (**B**) Selected cell cycle regulators were evaluated in the nine NF1-associated and one sporadic (STS26T) MPNST cell lines. (**C**) ST8814 and JH-2-002 transduced with doxycycline (Dox) inducible constructs expressing sh*PTPN11* #5003 or #818, were treated with vehicle or 300 ng/ml Dox for 72 hours, and RB phosphorylation at serine 780 or serine 807/811 was assessed. (**D**) ST8814 parental (par), or stable lines transduced with pLXSN-neomycin-E7 WT or E7 d21-24 mutant; and JH-2-002 parental, or stable lines transduced with pMSCV-puromycin-E7 WT or E7 d21-24 mutant, were treated with increasing dose of TNO155 for five days. Cell viability was evaluated by using the CCK-8 assay. (**E**) Cells as in D. were treated with DMSO or 0.3 μmol/L TNO155 for 48 hours.

To further explore signaling mechanisms that may potentiate TNO155 efficacy, we sought to identify effectors downstream of ERK signaling that may attenuate SHP2i dependence. It has been reported that the loss of RB1 function results in partial resistance to MEK and RAF inhibition in *BRAF*^V600E^ mutant melanoma cells (*33*), and given the high frequency of cell cycle dysregulation in MPNST, we focused on the RB tumor suppressor pathway. Notably, all MPNST cell lines on our panel have retained RB protein expression (Fig 3B), and none exhibited evidence of genomic alterations in *RB1* (Fig 2A), suggesting the presence of functional RB. RB1 plays a major role in the G1 cell cycle checkpoint which blocks S-phase entry (*34*), and phospho-RB (p-RB) is associated with RB1 inactivation. In *PTPN11*-depleted cell lines, p-RB was reduced, indicative of enhanced RB activity (Fig 3C), an observation also seen with SHP2 pharmacological inhibition with TNO155 in Figures 2D-F. These results demonstrate that ERK signaling inhibition via *PTPN11* genetic depletion, or treatment with SHP2i, induces RB hypo-phosphorylation, resulting in RB activation.

To evaluate if loss of RB1 function confers resistance to TNO155, we designed an experiment to inhibit RB1 function by introducing the ectopic expression of human papillomavirus (HPV) 16 E7. E7 inhibits RB1 function by binding to RB1, which disrupts the RB1/E2F complex leading to release and activation of E2F, and by enhancing RB1 degradation (*33, 35, 36*). While E2F activation by E7 can also lead to RB1 activation through p16 and CDK4/6 (*35*), this cannot occur in our panel of p16-null NF1-MPNST cell lines (Fig 3B). To test the hypothesis that LOF of RB1 confers TNO155 resistance, we generated stable cells expressing wild type (WT) E7 or E7 d21– 24 (a mutant incapable of binding to RB1), a dominant negative control. We observed that WT E7-expressing cells were more resistant to TNO155, relative to parental and E7 d21–24 mutant cells, with the IC50 increase of 5-fold in JH-2-002 and greater than 20-fold in ST8814 (Fig 3D). By measuring the direct protein expression of cyclin D1 and ERK signaling intermediates p-MEK and p-ERK, we found that treatment with TNO155 for 48 hours in E7 d21-24 mutant and parental cells was more potent than treatment in E7 WT cells (Fig 3E), suggesting that the E7 mutant with disabled RB1 binding restores sensitivity to TNO155. We therefore conclude that RB pathway is required for TNO155 sensitivity, and this evidence served as a justification for further exploration of targeting cell cycle/RB signaling in NF1-MPNST.

### Responses of NF1-MPNST cells to the CDK4/6i ribociclib

Inactivation of RB function as a consequence of recurrent loss of *CDKN2A* (encoding p16/ INK4a, Fig 3B), hyperactivation of CDK, and upregulated expression of *RABL6A* [a negative regulator of RB1, ref. (*27, 28*)] suggests that small-molecule CDK4/6i may be an additional therapeutic strategy in MPNST, particularly as a combination strategy. Indeed, 7/10 NF1-MPNST cell lines displayed limited sensitivity to the CDK4/6i ribociclib *in vitro*, by measuring metabolic activity as an indication of cell viability (Fig 4A). A complementary 3D microtissue assay revealed variable levels of sensitivity of tumor tissues derived from three individual MPNST PDX to ribociclib as well (Fig S4A). To understand mechanisms accounting for the insensitivity in some lines, we carried out a time-course combined with dose response study over 96 hours. We discovered that while all three CDK4/6i –abemaciclib, palbociclib, and ribociclib – effectively inhibited p-RB as a readout of reduction in Cyclin D-CDK4/6 complex activity in a dose-dependent manner, the three CDK4/6i all increased p-CDK4 at threonine 172 (T172), an indicator of CDK4 activation (Fig S4B), a phenomenon also recently observed in breast cancer cells which suggests that CDK4/6i stabilize the active p-T172 CDK4-cyclin D complex (*37–39*). We next tried to validate this observation and focused on ribociclib in an expansion cohort of eight NF1-MPNST cell lines (Fig 4B and Fig S4C). Treatment with 1μmol/L ribociclib effectively reduced p-RB and later phase effectors of cell cycle, PLK1 and cyclin A, but also caused a slight induction of ERK signaling, which in turn increased cyclin D1, cyclin E, and p-CDK4 T172 expression concurrently (Fig 4B and Fig S4C). This suggests that CDK4/6i induce an adaptive signaling response that reactivates the complex activity of cyclin D-CDK4/6, likely through the RTK/RAS/ERK pathway due to the high dependency of cyclin D expression on ERK signaling (*40*). To validate this hypothesis, we performed a proteomic profiling using human RTK phosphorylation array that detects 71 human RTK. Among these, tyrosine pan-phosphorylation of ErbB3, IGF-IR, Axl, and members in Ephrin signaling was up-regulated by ribociclib (Fig 4C and Fig S4D), which may further activate downstream RAS and cell cycle signaling, supporting the concept of combining CDK4/6i with agents targeting RAS/ERK pathway.

**Fig. 4.**
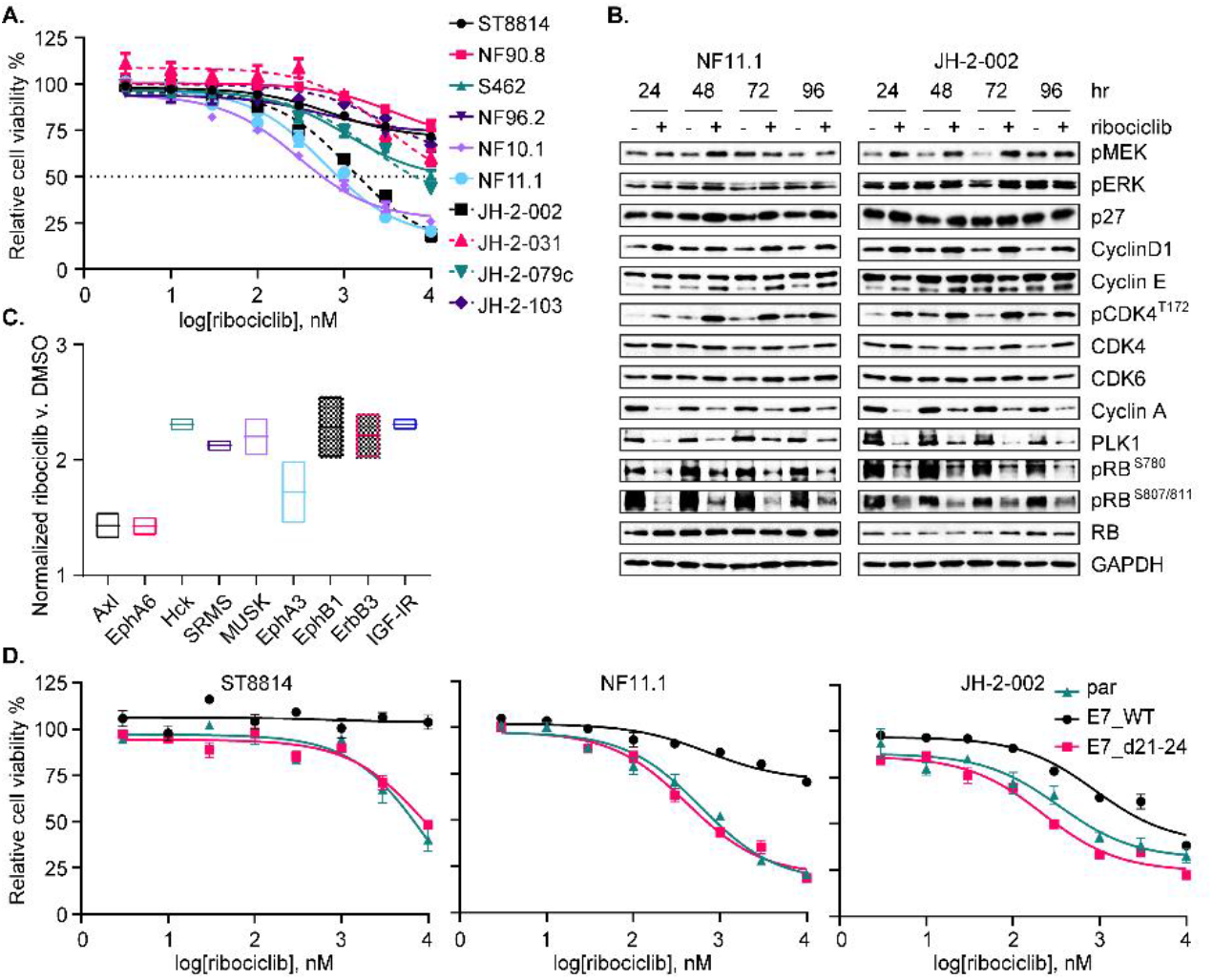
Responses of NF1-MPNST cells to the CDK4/6i ribociclib. (**A**) Ten NF1-associated MPNST cell lines were treated with increasing dose of the CDK4/6i ribociclib for five days. Cell viability was evaluated by using the CCK-8 assay. (**B**) Two NF1-MPNST cell lines were treated with DMSO or 1 μmol/L ribociclib over a time course. Signal intermediates in ERK and cell cycle pathways were assessed. (**C**) JH-2-002 cells were treated with DMSO or 1 μmol/L ribociclib for 24 hours. 71 phosphorylated human receptor tyrosine kinases (RTK) were evaluated using human RTK phosphorylation array C1. Signal intensity from technical duplicates was quantified using densitometry analysis, normalized to ribociclib v. DMSO, and significantly altered RTKs are shown. (**D**) Cells as in Fig 3E were treated with increasing doses of ribociclib for five days. Cell viability was evaluated by using the CCK-8 assay.

Previous reports revealed that in addition to the activation of upstream signaling pathways ERK/AKT/Hippo, the expression levels of selected cell cycle regulators (including CDK2/4/6, cyclins, p16, p21, E2F, p-CDK4 T172, RB, FOXM1, AURKA, p53 and MDM2/4), as well as the dissociation of p21 from cyclin D-CDK4 complex have putative relevance to CDK4/6i resistance (*38, 41*). We investigated additional explanations for CDK4/6i resistance observed in some cell lines. Specifically, we measured protein expression of several cell cycle regulators and signaling molecules in ERK/AKT pathways, but did not find a universal association of baseline expression of any single molecule with responsiveness to ribociclib across our NF1-MPNST cell line panel (Fig 3B and Fig S3A-B). As demonstrated in Figure 3 that RB function is critical for TNO155 efficacy, we also observed that it may be necessary for ribociclib efficacy as well, as resistance to ribociclib results from loss of RB function by WT E7, compared to mutant E7 d21-24 (Fig 4D). This suggests that CDK4/6 inhibition could be effective against MPNST in combination with SHP2i.

### Combined inhibition of SHP2 and CDK4/6 effectively suppresses MPNST cell growth

LOF of RB due to the selective upregulation of cell cycle signaling in MPNST relative to the precursor tumor, pNF, contributes partially to the insensitivity of NF1-MPNST to inhibitors of RAS/ERK signaling, including SHP2i and CDK4/6i (Fig 3 and 4). Therefore, we hypothesized that reinstatement of functional RB by CDK4/6i would enhance the treatment efficacy of SHP2i. In addition, SHP2i might attenuate the adaptive resistance to CDK4/6i through suppression of RTK/RAS/ERK signaling, thereby potentiating, and transforming the cytostatic effect of CDK4/6i to be cytotoxic. With this in mind, we tested the combinatorial effect of TNO155 plus ribociclib (referred to as the combination) on cell growth via three complementary methods and we found the additive activity of this combination against short- and long-term cell growth (Fig 5A-D and Fig S5A-B). Furthermore, the combination of high doses of TNO155 and ribociclib was able to overcome RTK-mediated trametinib resistance [Fig 5A and Fig S5B, ref. (*19*)]. To identify signaling changes that may lead to the additive effect, we investigated ERK and cell cycle pathways using eight NF1-MPNST cell lines following 72- and 96-hour treatment with TNO155 and/or ribociclib. Indeed, the combination mitigated ERK signaling (p-MEK and p-ERK), cyclin D1 expression, and p-CDK4 T172 induced by ribociclib, and more potently inhibited RB phosphorylation, E2F1, cyclin E/A and PLK1, in the majority of MPNST cell line models (Fig 5E and Fig S5C). Again, SHP2 knockdown by inducible sh*PTPN11* alleviated ribociclib-induced ERK signaling (Fig S5D), and resulted in greater decrease of cyclin A and PLK1 when combined with ribociclib (Fig 5F).

**Fig. 5.**
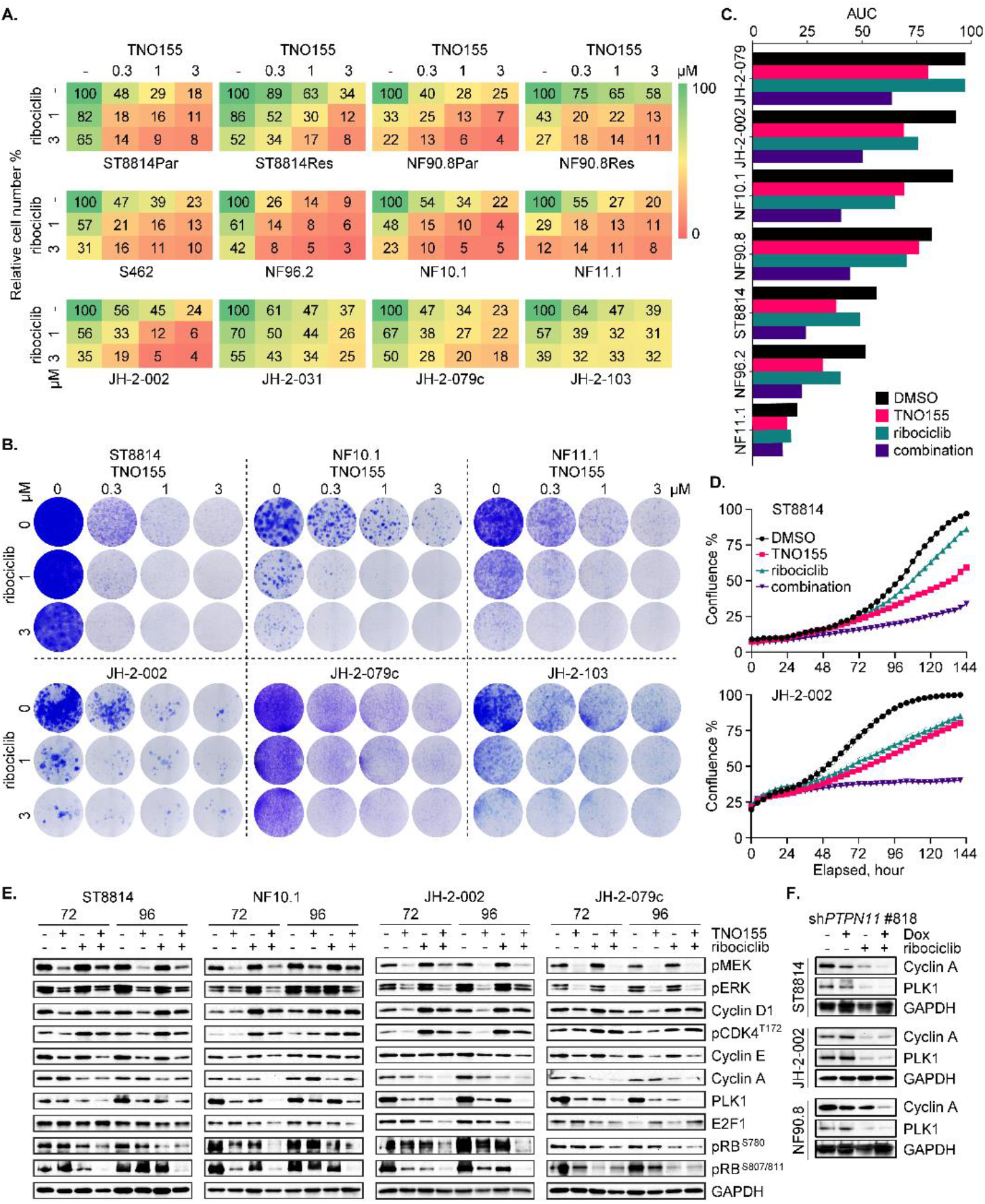
Combined inhibition of SHP2 and CDK4/6 effectively suppresses MPNST cell growth. (**A**) ST8814 and NF90.8 parental (Par) and trametinib-resistant (Res) lines, and eight NF1-MPNST cell lines were treated with DMSO, TNO155 (0.3, 1 and 3 μmol/L), ribociclib (1 and 3 μmol/L) and their combination for 7-10 days. Direct cell counting using trypan blue exclusion assay was performed by TC20 automated cell counter. (**B**) Cells as in A. were treated with drugs for 2-3 weeks, and colony formation was evaluated using crystal violet assay. (**C**) Area under the curve (AUC) is calculated based on the Incucyte cell confluence monitoring of the seven cell lines as shown, treated with DMSO, TNO155 (0.3 μmol/L), ribociclib (1 μmol/L) and their combination for six days. (**D**) Two NF1-MPNST cell lines were treated with DMSO, TNO155 (0.3 μmol/L), ribociclib (1 μmol/L) and their combination for six days. Cell confluence was monitored using Incucyte imaging systems. (**E**) Four NF1-MPNST cell lines were treated with DMSO, 0.3 μmol/L TNO155 and/or 1 μmol/L ribociclib for 72 and 96 hours. ERK signaling and cell cycle regulators were evaluated using immunoblot. (**F**) Three NF1-MPNST cell lines transduced with sh*PTPN11* #818 were pretreated with vehicle or 300 ng/ml Dox for 72 hours, followed by treatment with DMSO or 1 μmol/L ribociclib for additional 72 hours. Cell lysates were assessed for expression of the indicated proteins.

### Combination of TNO155 and ribociclib additively inhibits cell cycle and induces apoptosis

To deepen the understanding of mechanisms behind the increased efficacy of the combination, we performed RNAseq of JH-2-002 after treatment with TNO155 and/or ribociclib for 24 hours. The corresponding samples to RNAseq demonstrated signaling responses in line with our prior observations in Figure 5E (Fig S6A). As expected, 2503 genes were differentially expressed upon treatment with the combination compared to 1277 and 740 genes that were altered upon ribociclib and TNO155 treatment alone (Fig 6A). Furthermore, more than half of the genes differentially expressed upon treatment with the combination (1444 genes) were differentially expressed in the individual treatments (Fig 6A and Table S3). Pathway enrichment analysis against the KEGG database revealed that cell cycle signaling is significantly enriched in the intersection set ribociclib and the combination (Fig S6B). Deeper investigation into cell cycle signaling demonstrated an additive inhibition of mitotic prometaphase (Fig 6B) and cell cycle checkpoints (Fig 6C) by the combination, relative to TNO155 and ribociclib alone. To validate this finding, we carried out cell cycle distribution analysis in five NF1-MPNST cell lines (ST8814, NF10.1, JH-2-002, JH-2-079c and JH-2-103) as an aggregate using flow cytometry, and we saw an increasing trend of percent cells in G1 phase and a decreasing trend of percent cells in S phase when comparing cells treated with DMSO, TNO155, ribociclib, and the combination (Fig 6D).

**Fig. 6.**
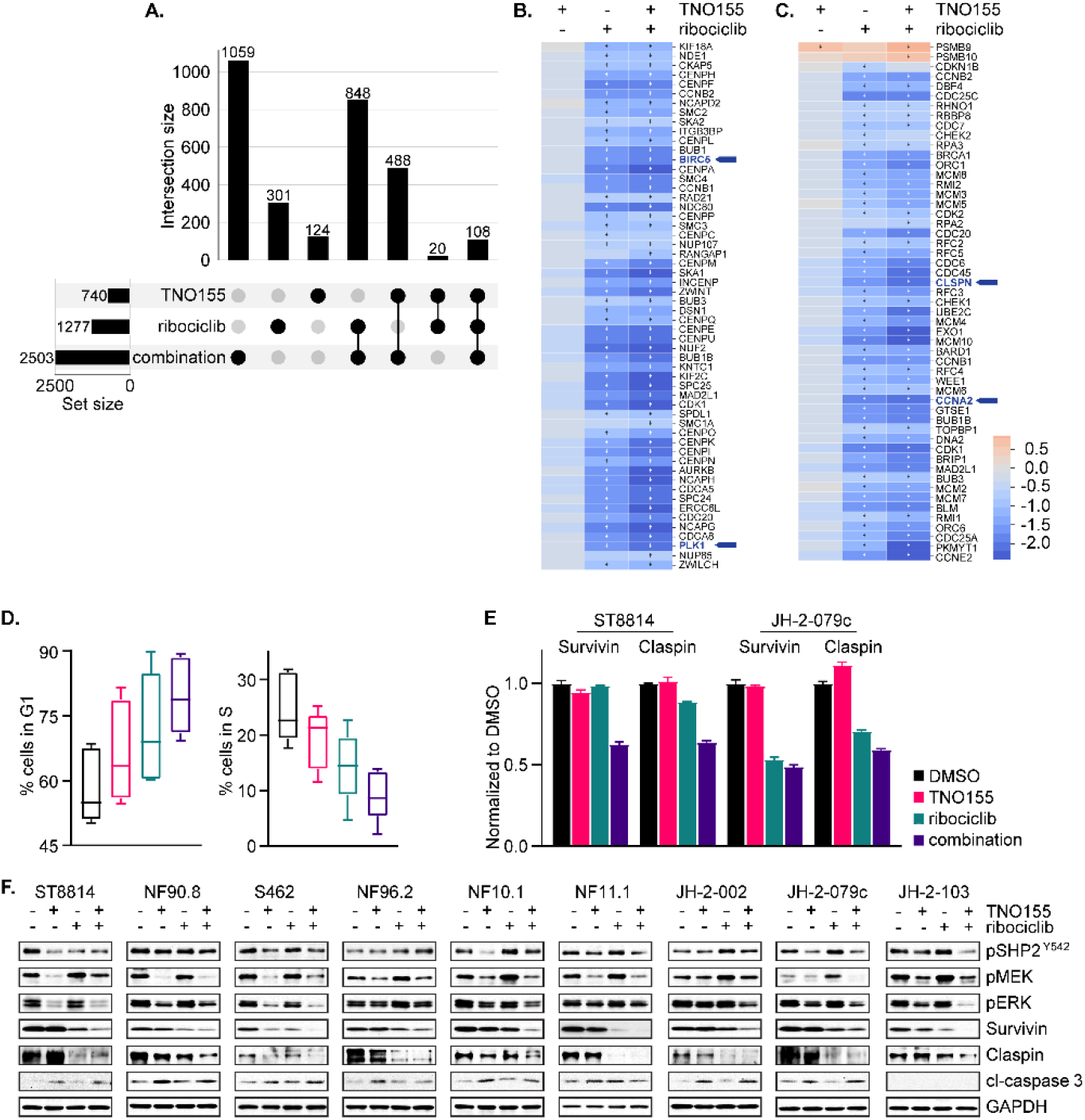
Combination of TNO155 and ribociclib additively inhibits cell cycle and induces apoptosis. (**A**) Upset matrix plot derived from RNAseq analysis, showing the overlapping numbers of significant genes (*P* adj< 0.05 and |FC|> 1.5) after 24-hour treatment with 0.3 μmol/L TNO155, 1 μmol/L ribociclib and their combination, normalized to DMSO control. (Rows=the sets; Columns=intersections between these sets). (**B** and **C**) Heatmaps demonstrating the more potent transcriptional inhibition of mitotic prometaphase (**B**) and cell cycle checkpoints (**C**) by combined TNO155+ribociclib relative to either single agent alone, highlighting *BIRC5*, *PLK1*, *CLSPN* and *CCNA2*. + = significance. (**D**) Five NF1-MPNST cell lines were treated with DMSO, 0.3 μmol/L TNO155 and/or 1 μmol/L ribociclib for 48 hours, following overnight starvation in 0.1% FBS-containing culture medium to synchronize cells. Cells were fixed in ice cold 70% ethanol and stained with propidium iodide/ RNase staining solution (Cell Signaling Technology, #4087), and then analyzed by flow cytometry. (**E**) ST8814 and JH-2-079c were treated with DMSO, 0.3 μmol/L TNO155 and/or 1 μmol/L ribociclib for 48 hours, and then protein lysates were assessed using human apoptosis antibody array (R&D systems, #ARY009). Signal intensity from technical duplicates was quantified by densitometry analysis using image J and normalized to DMSO. Data represent mean ± SEM. (**F**) Nine NF1-MPNST cell lines were treated as in E. and the indicated proteins involved in apoptosis and ERK signaling were detected using immunoblot.

As cell proliferation and programmed cell death are closely interconnected, we screened for the relative expression of 35 apoptosis-related proteins under these four conditions. Using one traditional and one patient-derived cell lines as a representative cohort, we found universally decreased expression of two inhibitors of apoptosis proteins (IAP) survivin/*BIRC5* and claspin/*CLSPN* in ST8814 and JH-2-079c treated with the combination (Fig 6E and Fig S6C), and increased caspase-3/7 activation (Fig S6D). *survivin/BIRC5*, *claspin/CLSPN*, *PLK1*, and *CCNA2* are also down-regulated upon the combination treatment (Fig 6B-C, highlighted), suggesting that the regulation occurs at the transcriptional level. In an expanded validation cohort of nine NF1-MPNST cell lines, survivin, claspin, and PLK1 were more potently inhibited by the combination, relative to either single agent alone (Fig 6F and Fig S6E). These results demonstrate that the combined use of TNO155 and ribociclib prevents signaling adaptation to CDK4/6i and converges on the restoration of RB function, which in turn induces G1 cell cycle arrest and transcriptionally inhibits the IAP survivin and claspin to activate caspase-3/7, subsequently activating apoptosis.

### Combination of TNO155 and ribociclib is active against MPNST tumor growth *in vivo*

Given our results demonstrating the additive *in vitro* effects of the combination, we hypothesized that SHP2i plus CDK4/6i may be the superior combination of RAS signaling pathway inhibitors in MPNST, due to better or comparable *in vivo* efficacy compared to SHP2i plus MEKi, and have more favorable tolerability, as judged by mouse body weight in both immuno-compromised and competent models (*42–45*). To address whether this combination is active in *in vivo* models of MPNST, we utilized a total of six unique PDX models to assess the combination versus either single agent, using doses (TNO155 7.5 mg/kg, twice daily; ribociclib 75 mg/kg, once daily) lower than those selected in tolerability studies (*43*). In fact, TNO155 has surprisingly potent single agent activity across the six PDX over 4 weeks (Fig 7A and B), compared to our previous results using the tool compound SHP099 (*20*). We thus defined the top three PDX that were almost equally sensitive to the combination and TNO155 alone as very sensitive (VS, Fig 7A), and the bottom three that displayed differential sensitivity to the combination and TNO155 as partially sensitive (PS, Fig 7B). We directly compared the efficacy of TNO155 alone and TNO155/ ribociclib combination against the TNO155/ MEKi combination. We observed that the most potent anti-tumor effects were seen with TNO155+ ribociclib, relative to TNO155 or TNO155+ trametinib, across a panel of five PDX (Fig S7A), consistent with previous studies in both RTK-driven and *KRAS*-mutant cancers (*43*).

**Fig. 7.**
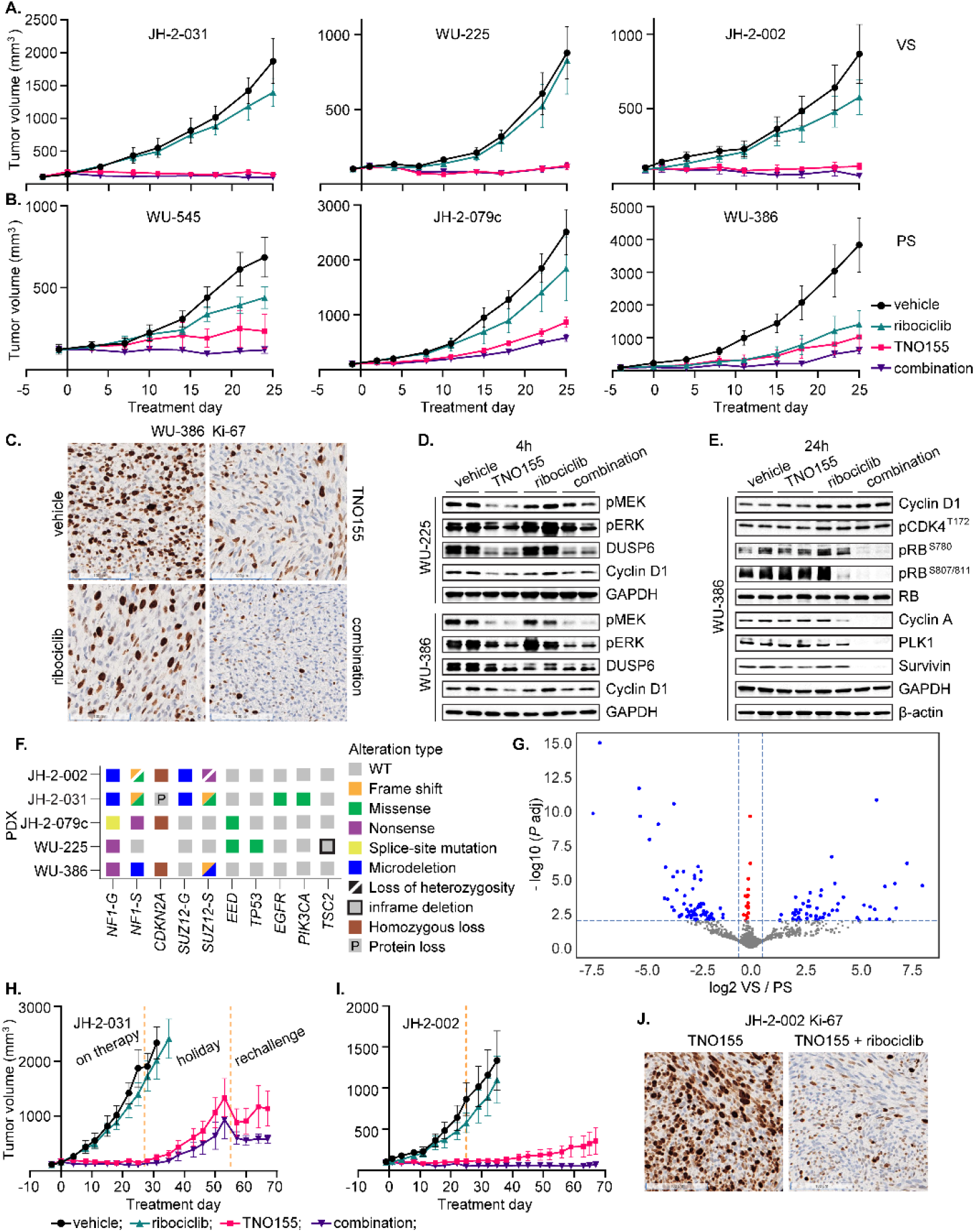
The combination of TNO155 and ribociclib is active against MPNST tumor growth *in vivo*. (**A**-**B**) 5- to 6-week-old female NRG mice bearing six individual NF1-MPNST patient-derived xenografts (PDX) were treated with vehicle, ribociclib (75 mg/kg, once daily), TNO155 (7.5 mg/kg, twice daily) or their combination by oral gavage for about four weeks. Tumor size was measured twice weekly. Average tumor volume of 4-5 mice per arm was plotted over the time course of treatment days. VS= very sensitive; PS= partially sensitive. (**C**) Tumors of each arm from the PDX WU-386 were harvested four hours post last dose of 4 weeks on drugs and fixed in 10% NBF. The Ki-67 expression was assessed using IHC. (**D**-**E**) Tumors of each arm from the PDX WU-225 and WU-386 were harvested 4 hours post last dose, 4 weeks on drugs (**D**), or 24 hours post last dose, 3 days on drugs (**E**). The indicated proteins involved in ERK and cell cycle signaling were detected by immunoblot. (**F**). Onco-print of key driver genes (*NF1, CDKN2A, SUZ12* and *EED*) in MPNST and putative others across a panel of MPNST PDX is shown. For *NF1* and *SUZ12*, both germline (G) and somatic (S) mutations are shown. (**G**) Volcano plot demonstrating log2 fold change of VS/PS as a function of −log10 (*P* adj). Blue dots represent 227 genes that were significantly altered (*P* adj< 0.05 and |L2FC| > 0.585) when comparing the RNA seq data of VS (JH-2-031, WU-225 and JH-2-002) v. PS (WU-545, JH-2-079c and WU-386). (**H**) The mice bearing the PDX JH-2-031 were on initial treatment as in A. for about four weeks, and then left untreated for another four weeks before rechallenging with the combination for additional two weeks. (**I**) The mice bearing the PDX JH-2-002 as in A. were continuously on treatment for ten weeks. (**J**) Tumors of each arm from the PDX JH-2-002 were harvested four hours post last dose of 10 weeks on drugs and fixed in 10% NBF. The Ki-67 expression was assessed using IHC.

In addition, we investigated the pharmacodynamic (PD) activity of the combination to identify potential molecular response features underlying the therapeutic response. We biochemically characterized ERK and cell cycle signaling at 4 hours post 4-week treatment (Fig 7C-D and Fig S7B-C), and 24 and 48 hours after 3-day treatment (Fig 7E and Fig S7D). Ki-67 expression, a marker of proliferation, was more potently reduced by the combination compared to TNO155 or ribociclib, as evidenced by immunohistochemistry (IHC) staining at the end points of treatment (Fig 7C, and Fig S7B). Early and end-point *in vivo* PD studies were in agreement with what we saw *in vitro* that the combination relieved the adaptive response to ribociclib and led to greater suppression of cell cycle regulators, such as p-RB, cyclin A, PLK1 and survivin (Fig 7D-E and Fig S7C-D).

To ask whether genomic or expression biomarkers exist to predict *de novo* sensitivity to TNO155, we next analyzed whole exome sequencing (Fig 7F) and RNAseq data (Fig 7G) derived from VS PDX (JH-2-002, JH-2-031 and WU-225) and PS PDX (JH-2-079c, WU-545 and WU-386). The VS PDX JH-2-031 with activating *PIK3CA* Q546K mutation and WU-225 with *TSC2* in frame deletion retained sensitivity to TNO155 (Fig 7F). Several putative molecules relevant to drug resistance, such as WNT2B and NTRK2 were significantly upregulated (*P* adj<0.05 and |fold change|>1.5) in PS compared to VS models (Fig 7G and Fig S7E, Table S4). Although we did not observe a difference in sensitivity between TNO155 alone or TNO155 plus ribociclib at 4 weeks of treatment in the VS PDX, there is a sufficient concern that single-agent treatment in this aggressive malignancy would be insufficient for anything longer than an immediate short-term response. To investigate this further, we retreated the PDX JH-2-031 cohorts of TNO155 single agent and the combination with TNO155 plus ribociclib after a drug holiday (Fig 7H), or treated the PDX JH-2-002 for an extended course with TNO155 alone or the combination (Fig 7I). We observed combination benefit regardless of intermittent (Fig 7H) or continuous dosing of therapy over a 10-week treatment (Fig 7I), indicating that the combination resulted in more profound and/or more durable anti-tumor effects which was supported by a significant difference in Ki-67 (Fig 7J), and that treatment-emergent resistance to TNO155 may limit the duration of response.

## DISCUSSION

Our collective data demonstrate that cell proliferation and tumor growth of NF1-MPNST are SHP2/*PTPN11*-dependent. Using both SHP2 genetic ablation and pharmacological inhibition we observe effective reduction in MPNST cell proliferation through ERK pathways and downstream cell cycle suppression. Reconstitution of NF1 GAP function mediates resistance to TNO155, suggesting that NF1-null MPNST are sensitive to SHP2i and further that NF1 loss of function may be an important genomic biomarker for patient selection as clinical trials of SHP2i advance. TNO155 treatment causes a transcriptional response which significantly overlaps with that induced by MEKi, suggesting that the effects of SHP2i are largely mediated through ERK, although other RAS effector pathways may exert secondary roles. This redundancy in pathway inhibition suggests the potential for overlapping clinical toxicity and so alternate combination strategies that enhance the anti-tumor effects of SHP2i and maintain tolerability may be important for trial design.

Consistent with previous studies (*28*), cell cycle signaling is significantly upregulated in MPNST v. pNF, and results in LOF of RB, suggesting the role for CDK4/6i as part of a combination strategy. Interestingly, NF1-MPNST cells are insensitive to CDK4/6i, likely at least in part due to adaptive resistance resulting from induction of RTK/ERK signaling and reactivation of cyclin D-CDK4/6 complex activity. While we did not find any correlation between baseline expression markers and sensitivity to TNO155 and/or ribociclib, our data suggest that functional RB is critical for both TNO155 and ribociclib sensitivity. TNO155 plus ribociclib activity converges on restoration of biologically active RB, otherwise inactivated by upregulation of CDK and downregulation of p16, and on greater inhibition of later-phase cell cycle regulators PLK1 and cyclin A. Mechanisms of anti-proliferation and anti-tumor effects involve an additive impact on global transcription as well as an additive repression in gene transcription related to cell cycle signaling. The combination leads to greater G1 cell cycle arrest, and also more potently inhibits two apoptotic regulators, survivin and claspin, and induces downstream caspase activation to promote apoptosis.

Lastly, we assessed a total of six PDX models for the efficacy of TNO155, ribociclib, and their combination. We found that three PDX have a near complete response to TNO155 single agent with no apparent difference induced by the combination in the short-term, while the other three are partially sensitive and have more complete responses with the addition of ribociclib. Overall, the combination is strikingly active in all *in vivo* PDX models of NF1-MPNST tested. RNAseq analysis of the MPNST PDX revealed differential expression of multiple putative resistance-relevant genes involved in Wnt/β-catenin/YAP/TAZ signaling pathways in the three PS models, suggesting that these pathways may mediate resistance to TNO155, and agents targeting these pathways may be another potential viable partner with SHP2i, which remains to be explored.

Clinical trial efforts have focused on dysregulated RAS effector pathways that are critical to MPNST tumorigenesis, but to date have been unsuccessful in people with advanced MPNST. This study suggests that despite past failures using downstream inhibitors of RAS signaling, SHP2 may in fact represent a critical node to inhibit MPNST tumorigenesis due to the loss of NF1 and that combinations of SHP2i with inhibitors of other critical effectors could have greater success in the clinic. Our extensive preclinical data demonstrate that clinically available inhibitors of SHP2 and CDK4/6 in combination and at well-tolerated dosing regimens result in deep and durable pre-clinical, anti-tumor responses in mouse models with similar efficacy as MEKi plus SHP2i. In addition, this combination is already under investigation in an ongoing trial (NCT04000529) and data to support the clinical safety profile of TNO155 plus ribociclib are expected to result soon. Our evidence is in favor of the advancement of novel SHP2i into clinical trials for the treatment of patients with MPNST.

Both SHP2i and CDK4i are considered pro-senescence cancer therapies, which disable cell proliferative capacity and trigger an inflammatory process that ultimately eliminates the tumor cells (*49*). Here we have provided a large body of evidence demonstrating the synthetic lethality of TNO155 plus ribociclib in both clinically relevant *in vitro* and *in vivo* models of MPNST. However, these models are limited to the tumor-cell-intrinsic signaling response and utilize immunocompromised mice, which preclude the evaluation of the combination on anti-tumor immunity. As such, further studies on treatment efficacy and the role of the combination in modulation of tumor immune microenvironment are needed in immunocompetent and humanized mouse models of MPNST, aiming to identify novel therapeutic opportunities based upon mechanistic rationale.

We have generated strong pre-clinical evidence in support of this combination, suggesting that a clinical study evaluating the efficacy and tolerability of TNO155, alone or in combination with ribociclib, should be pursued in patients with MPNST. Future studies will identify potential genomic or expression biomarkers that predict treatment efficacy to facilitate selection of patients who will have clinical benefit. With the clinical testing of TNO155 and ribociclib currently underway in patients with other solid tumors, these studies are timely, relevant, and will serve as an important contribution to the advancement of novel SHP2i combinations in clinical use. Finally, acquired resistance to targeted therapy is common and inevitable. Exploration of resistance mechanisms will identify novel combination partners with SHP2i and move SHP2i-centric therapeutics forward for optimal clinical application.

## MATERIALS AND METHODS

### Study design

We recently reported that the adaptive response to MEKi in MPNST involves the upregulation of activity of multiple RTK (*19, 20*). We therefore hypothesized that SHP2 inhibition might be a strategy to overcome the global RTK upregulation seen with single agent MEKi, and we subsequently demonstrated that combined MEKi/ SHP2i is additive in both *in vitro* and *in vivo* models of NF1-MPNST (*20*). Here, we focus on SHP2i as a primary strategy to inhibit MPNST proliferation and tumor growth and identify CDK4/6i as a potential partner agent to be tested in combination with SHP2i, thus avoiding the overlapping toxicity of two agents that both inhibit phospho-ERK. Conversely, as loss of *CDKN2A* is a common feature in MPNST, we hypothesized that CDK4/6i may be active and further potentiated by drugs targeting upstream regulators of RAS, such as SHP2. *In vitro* experiments were performed at least two times. For *in vivo* studies, five mice per arm were randomly assigned to either agent alone, or the combination, and followed for at least 4 weeks. The investigators were not blinded to group allocation during data collection and analysis.

### Cell lines, antibodies, and reagents

Four patient-derived NF1-MPNST cell lines (JH-2-002, JH-2-031, JH-2-079c and JH-2-103) and eight PDX [JH-2-002, JH-2-031, JH-2-079c, WU-225, WU-386, WU-545, MN-2 and MN-3-002 (the last two for 3D tumor microtissue)] were generated in our laboratories at Johns Hopkins (JH), Washington University (WU, St. Louis) or University of Minnesota (MN) from biospecimens collected during surgical resection from patients with NF1 (*8, 30*). Material was collected under Institutional Review Board–approved protocols (JHU #J1649; WU #201203042; and MN #STUDY00004719). All patients provided written informed consent. All cell lines used in these experiments were verified by short-tandem repeat (STR) profiling for cell line authentication at Johns Hopkins University Core Facility, tested negative for Mycoplasma contamination, and passaged *in vitro* for fewer than 3 months after resuscitation. All growth media were supplemented with 10% FBS, 2 mmol/L l-glutamine, and 1% penicillin–streptomycin. Trametinib-resistant cell lines were maintained in complete growth medium plus 20 nmol/L of trametinib (*19*).

TNO155, ribociclib, and trametinib were provided under a Materials Transfer Agreement with Novartis Institute for Biomedical Research (NIBR). Palbociclib, abemaciclib, and SHP099 were purchased from SelleckChem. Drugs for *in vitro* studies were dissolved in DMSO to yield 10 or 1 mmol/L stock solutions, and stored at −20°C.

The sources and catalog numbers of cell lines, plasmids, antibodies, and reagents are detailed in Table S5.

### Immunoblotting

Cells were disrupted on ice in RIPA lysis buffer. Protein concentration was determined with Pierce BCA protein assay kit (#23227, Thermo Fisher Scientific). Equal amounts of protein were separated by SDS-PAGE, transferred to nitrocellulose membranes, immunoblotted with specific primary and secondary antibodies, and detected by chemiluminescence with the ECL detection reagents, Immobilon Western chemiluminescent HRP substrate (#WBKLS0500, Millipore), or Pierce ECL Western blotting substrate (#32106, Thermo Fisher Scientific). The membranes were imaged using ChemiDoc touch imaging system (Bio-Rad).

### Cell proliferation assay

Cells were seeded in 96-well plates at 2,000 cells per well. A dose range of the compound indicated was prepared by serial dilutions and then added to the dishes containing adherent cells. Cells were incubated with drug for the indicated time. Cell growth was quantitated using the Cell Counting Kit-8 (Dojindo). For each condition, three replicates of each concentration were measured. Relative survival in the presence of drugs was normalized to the untreated controls after background subtraction. Graphs were generated using Prism 9 based on the average of three replicates.

### Active RAS pull-down assay

Cells were seeded in 10-cm dishes. The following day, the 70% to 80% confluent cells were collected, and GTP-bound RAS was quantified using active RAS detection kit (# 8821) from Cell Signaling Technology according to the manufacturer’s instructions.

### Lentivirus-based inducible shRNA-mediated knockdown cells

shRNAs targeting *PTPN11* #5003 and #818 [oligo sequences listed in Table S6, ref. (*24*)] were subcloned into Tet-pLKO-puro vector (Addgene #21915). The lentiviruses encoding sh*PTPN11* were packaged in HEK293T cells. The target cells were infected overnight by filtered viral supernatant in the presence of 8 μg/mL of polybrene (#TR-1003-G, Millipore). The infected cells were selected with puromycin (2 μg/mL) for one week before the analysis of the knockdown effects.

### Retrovirus-based gene expression cell system

The human *NF1-GRD* (*46*) was amplified using the gDNA from HEK293T as PCR template and sub-cloned into TTIGFP-MLUEX vector harboring Tet-regulated promoter (Primer sequences are listed in Table S6). The retroviruses encoding the rtTA3, NF1-GRD or E7, E7 d21-24 were packaged in Phoenix-AMPHO cells. The medium containing virus was filtered with 0.45μm PVDF filters followed by incubation with the target cells for 8 hours. The cells were cultured in virus free medium for two days and then selected with hygromycin (300 μg/ml), puromycin (2 μg/ml) or neomycin (800 μg/ml) for one week. For TTIGFP-MLUEX doxycycline inducible system, the positively infected cell populations were further sorted using FACSMelody cell sorter (BD) with GFP as a marker after overnight exposure to 1 μg/ml doxycycline and the sorted positive cells were cultured and expanded in medium without doxycycline but with antibiotics at a maintaining dose until the following assays.

### Colony formation assay

NF1-MPNST cells were treated with DMSO, TNO155 (0.3, 1 and 3 μmol/L), and/or ribociclib (1 and 3 μmol/L) for 2-3 weeks. Cells were washed with PBS, fixed with 10% neutral buffered formalin, and then stained with 0.1% crystal violet for 30 minutes.

### phospho-RTK array

Proteome profiler human RTK phosphorylation array C1 (#AAH-PRTK-1, Ray Biotech) was used to evaluate 71 p-RTK as per manufacturer instructions. Briefly, cells were rinsed with cold PBS and lysed in lysis buffer, 500 μg lysates were incubated with blocked membranes overnight at 4°C. Membranes were subsequently washed, incubated with diluted HRP detection antibody and chemiluminescent reagent mix, and imaged on the ChemiDoc touch imaging system.

### Apoptosis antibody array

NF1-MPNST cells were treated with DMSO, 0.3 μmol/L TNO155, and/or 1 μmol/L ribociclib for 48 hours and then 500 μg protein lysates were assessed using human apoptosis antibody array (#ARY009, R&D systems).

### Cell cycle analysis by flow cytometry

Modulation of the cell cycle was determined in NF1-MPNST cells treated with DMSO, TNO155, ribociclib, or their combination for 48 hours, following incubation with culture medium containing 0.1% FBS for 24 hours to synchronize cells. After trypsinization, cells were fixed in ice-cold 70% ethanol for at least 30 minutes and were then labeled with propidium iodide (PI)/RNase staining solution (#4087, Cell Signaling Technology) and further incubated for 15 minutes at 37°C. Finally, cells were analyzed using the FACSCelesta™ Cell Analyzer (BD Biosciences). Data analysis was performed using FlowJo 10.8, and cell cycle distribution was assigned by using the implemented models.

### *In vivo* mouse studies

NOD Rag gamma (NRG, #007799) female mice were purchased from the Jackson Laboratory. All mouse experiments were approved by the Institutional Animal Care and Use Committee (IACUC) at Johns Hopkins. Minced tumor fragments from donor mice were implanted subcutaneously close to the sciatic nerves of 6- to 8-week-old NRG female mice. Drug treatment was started when tumor size reached roughly 100-200 mm^3^. Mice were randomized into treatment groups by an algorithm that arranges animals to achieve the best-case distribution to ensure that each treatment group has similar mean tumor burden and SD. Vehicle, trametinib (0.15 mg/kg for JH-2-079c and 0.075 mg/kg for others, once daily, Novartis; dissolved in 0.5% hydroxypropyl methyl cellulose and 0.2% Tween 80 in water, pH 8), TNO155 (7.5 mg/kg, twice daily, Novartis; dissolved in 0.5% methyl cellulose and 0.1% Tween 80 in water), ribociclib (75 mg/kg, once daily, Novartis; dissolved in 0.5% methyl cellulose in water), or their combinations were administrated by oral gavage, based on mean group body weight, with treatment schedule of 5 days on/2 days off. The endpoint of the experiment for efficacy studies was considered 4 weeks on treatment (extended based on depth of response) or the longest tumor diameter of 2 cm as per our approved protocol, whichever occurred first. Tumors were measured twice weekly by calipering in two dimensions, and tumor volume was calculated by: L × W^2^(π/6), where L is the longest diameter and W is the width.

### Whole exome sequencing (WES)

WES FastQ files were trimmed by using Trimmomatic v 0.39 (*47*) and aligned against reference sequence hg38 via BWA-MEM (*48*). Duplicated reads were marked by using PICARD MarkDuplicates function. GATK V4.2 base quality score recalibration (BQSR) was also used to process BAM files. For PDX sequence data, Xenosplit was used to filter out mouse-derived reads using mouse (GRCm38) and human (hg38) reference genomes. Somatic SNVs and small indels were detected using VarScan2 (*49*), Strelka2 (*50*), MuTect2 (*51*), and Pindel (*52*). Variant filtering and annotation were done by using Variant Effect Predictor (VEP) (*53*). Common variants found in the 1000 Genomes MAF and GnomAD MAF > 0.05 were filtered out.

For *NF1*, *SUZ12* and *CDKN2A* germline and somatic deletion, BAM files from normal blood or tumor were used for the germline or somatic deletion calling respectively by using GATK 4.2. The main steps included: 1. CollectReadCounts function was used to extract the counts of reads based on the IDT xGen Exome Research Panel v1.0 BED file; 2. DetermineGermlineContigPloidy function was used to determine the baseline contig ploidy for germline samples given the counts data; 3. GermlineCNVCaller was used to call copy-number variants in germline samples; 4. BAF (the B-Allele Frequency) was called by Hatchet 1.2.1, and normalized read ratio of tumor to normal, and BAF data were used to call somatic gene deletion.

### Bulk RNA-seq library preparation, sequencing, and analysis

Primary tumor samples derived from 7 MPNST v. 31 pNF (Fig 3A), PDX samples from six patients (Fig 7G) and 30 samples (triplicate in 10 conditions) from JH-2-002 cells were used for RNAseq. The library was prepared using TrueSeq stranded total RNA kit with Ribo-Zero for rRNA depletion and was sequenced by NovaSeq6000 S4 (150bp PE) with targeted coverage of 60M total reads per sample.

FastQ files were aligned to GRCh38. RNA reads were quantified using the Salmon algorithm (*54*). Raw counts were normalized, and the DESeq2 package was used to call the differentially expressed genes (*55*). Exploratory data analysis was performed using the MAGINE software package, and ggplot2 was applied to draw the volcano plots (*56*).

### Incucyte cell proliferation assays

2,000 cells per well were plated in 96-well plates and treated with the indicated dose of drugs or caspase-3/7 dye for apoptosis (Sartorius, #4440). Cell proliferation and apoptosis were monitored by a microscope gantry that was connected to a network external controller hard drive that gathered and processed image data, using the IncuCyte Live-Cell Imaging System (Essen BioSciences, Ann Arbor, MI), which provides a time-lapse and an automated in-incubator method for quantifying cell growth.

### Immunohistochemistry (IHC)

Ki-67 IHC was performed at Johns Hopkins IHC core facility. Briefly, immunolabeling for Ki-67 was performed on formalin-fixed, paraffin-embedded sections on a Ventana Discovery Ultra autostainer (Roche Diagnostics). Following dewaxing and rehydration on board, epitope retrieval was performed using Ventana Ultra CC1 buffer (#6414575001, Roche Diagnostics) at 96°C for 48 minutes. Primary antibody anti-Ki-67 (1:200 dilution; #Ab16667, lot number GR3185488-1, Abcam) was applied at 36°C for 60 minutes. Primary antibodies were detected using an anti-rabbit HQ detection system (#7017936001 and 7017812001, Roche Diagnostics) followed by Chromomap DAB IHC detection kit (#5266645001, Roche Diagnostics), counterstaining with Mayer’s hematoxylin, dehydration, and mounting. Whole slide imaging was performed at the Oncology Tissue Services Core of Johns Hopkins University. Scanning was carried out at up to 40X magnification (0.23 microns/pixel) using a Hamamatsu Nanozoomer S210 digital slide scanner (Hamamatsu Photonics, Shizuoka, Japan). Whole slide images were visualized in Concentriq digital pathology platform (Proscia, Philadelphia, PA), and representative images at 20X magnification were shown.

### 3D tumor microtissue

3D microtissue assay was described previously (*9*). All mouse experiments were approved by the IACUC at University of Minnesota under protocol #2101-38758A. For each PDX, a sample was minced in a basement membrane matrix and passed through a 1mL syringe (18-gauge needle) and injected into the flank of NRG mice. Once the xenograft tumors reached the maximum size allowed (2000 mm^3^), mice were euthanized, tumors were extracted in a laminar flow hood under sterile conditions and were immediately digested using the human tumor dissociation kit (130-095-929, Miltenyi Biotec) in combination with the GentleMACS Octo Dissociator (Miltenyi Biotec). The digested PDX was filtered through a 70 μm cell strainer and then depleted of residual red blood cells using 1X RBC Lysis Buffer (eBioscience). Dead cells were removed from the digested PDX using a dead cell removal kit (130-090-101, Miltenyi Biotec). Murine cells were removed from the dead-cell-depleted fraction using a mouse cell depletion kit (130-104-694, Miltenyi Biotec). Cell viability was determined using flow cytometry and 7-AAD viability staining solution (Biolegend). Cell counts were ascertained using the Countess (Invitrogen).

Microwell plates were fabricated following previously established methods (*57*). For the assembly of collagen microtissues, previously established protocols were followed (*57, 58*). Briefly, high-concentration rat tail collagen I (Corning) was buffered with 10 × Dulbecco’s phosphate-buffered saline (DPBS), neutralized to pH 7.4, supplemented with 10% Matrigel (Corning), diluted to 6 mg/mL concentration, and mixed with cells (6×10^6^ cells/mL). At 4°C, the collagen solution was partitioned into droplets using a flow-focusing microfluidic device. Tissues were collected in a low-retention Eppendorf tube and polymerized for 30 min at 25°C. The oil phase was then removed and the collagen microtissues were resuspended in culture medium.

Microtissues containing encapsulated PDX cells were fabricated and cultured. To assess cell viability, constructs were washed thoroughly with DPBS (Corning) and then incubated in the dark at 21°C for 30 minutes with a staining solution of 5 μM DRAQ5 (Invitrogen) and 5 μM Calcein AM (Thermo Fisher). A Zeiss Axio Observer was used to image z-positions at 50 μm intervals over a distance of 250 μm and images were further analyzed using Cellpose (*59*) and ImageJ (NIH). For experimental wells, live cell over total cell count ratio was normalized to vehicle-only treated cells to produce percent cell viability. Data were analyzed with GraphPad Prism 9 and dose-response curves generated using nonlinear regression Log(inhibitor) vs. response-variable slope model.

### Statistical analysis

Two-way ANOVA was used to calculate statistical significance. Analyses were considered statistically significant if adjusted *P* < 0.05.

## Supplementary Materials

This PDF file includes:

Figures S1 to S7 and related figure legends\

Data files Table S1 to S6 (Excel files)

## Acknowledgments

This work was partially supported by the NF Research Initiative (NFRI) which was made possible by an anonymous philanthropic gift to the Multidisciplinary Neurofibromatosis Program at Boston Children’s Hospital.

We are grateful to Drs. Gregory Riggins, Margaret Wallace, and Angelina Vaseva for providing cell lines; to Sujayita Roy of the Oncology Tissue Services for assistance with Ki-67 IHC; to Dr. Kyle B. Williams and Justin F. Tibbitts for microtissue technical assistance and discussions; and to Jennifer Meyers and Dixie Hoyle of the Experimental and Computational Genomics Core for assistance with RNA-seq (Core Facilities of SKCCC at Johns Hopkins).

## Funding

This work was supported by grants from:

Novartis Institute for Biomedical Research (CAP)

The NF Research Initiative (ACH, SJG, DL, DKW, and CAP)

Hyundai Hope on Wheels (CAP)

the Neurofibromatosis Therapeutic Acceleration Program (CAP)

the Children’s Cancer Foundation (CAP)

the SKCCC Cancer Center Core NIH P30 CA006973

## Author contributions

Conceptualization: JW, CAP

Methodology: JW, AC, LZ, JCP, YL, KP, XZ, ATL, EC, NL, DKW, DAL, SEM, SJG, ACH, CAP

Investigation: JW, AC, LZ, JCP, YL, KP, XZ, ATL, EC, NL, DKW, DAL, SEM, SJG, ACH, CAP

Visualization: JW, AC, LZ, JCP, YL, ATL, NL, SJG

Funding acquisition: CAP

Project administration: SEM, CAP

Supervision: CAP

Writing – original draft: JW, SJG, CAP

Writing – review & editing: JW, AC, LZ, JCP, YL, KP, XZ, ATL, EC, NL, DKW, DAL, SEM, SJG, ACH, CAP

## Data and materials availability

RNAseq data are deposited in the Sage Bionetworks NF Data Portal (https://www.synapse.org) with the identification number syn35589856. All other data are available in the main text or the supplementary materials.

## Abbreviations

3D: Three dimensional
60M: 60 million
7-AAD: 7-aminoactinomycin D
adj: adjusted
AKT: Ak strain transforming
ANOVA: Analysis of variance
AS: Anti-sense
AUC: Area under the curve
AURKA: Aurora kinase A
BAF: B-allele frequency
BAM: Binary Alignment Map
BCA: Bicinchoninic acid
BD: Becton, Dickinson and Company
BED: Browser Extensible Data
BIRC5: Baculoviral IAP repeat containing 5
BQSR: Base quality score recalibration
BWA-MEM: Burrows-Wheeler Alignment Tool - Maximal Exact Match
°C: Degree Celsius
CC1 buffer: Cell Conditioning buffer
CCK-8: Cell Counting Kit-8
CCNA2: Cyclin A2
CDK: Cyclin dependent kinase
CDK4/6i: CDK4/6 inhibitor
CDKN2A: Cyclin dependent kinase inhibitor 2A
CLSPN: Claspin
cm: Centimeter
ctrl: Control
DAB: 3,3′-Diaminobenzidine
DEG: Differentially expressed genes
DMSO: Dimethyl sulfoxide
Dox: Doxycycline
DPBS: Dulbecco’s phosphate-buffered saline
ECL: Enhanced chemiluminescence
EED: Embryonic Ectoderm Development
ERBB3: Erb-b2 receptor tyrosine kinase 3
ERK: Extracellular signal-regulated kinase
F: Forward
FACS: Fluorescence-Activated Cell Sorting
FBS: Fetal bovine serum
FOXM1: Forkhead box M1
G: Germline
GAP: GTPase-activating protein
GATK: Genome Analysis Toolkit
gDNA: genomic deoxyribonucleic acid
GEF: Guanine nucleotide exchange factors
GFP: Green fluorescent protein
ggplot2: Grammar of graphics plot 2
GnomAD: Genome Aggregation Database
GRCh38: Genome Reference Consortium Human Build 38
GRD: GAP related domain
GTP: Guanosine triphosphate
HEK: Human embryonic kidney
HPV: Human papillomavirus
HRP: Horseradish peroxidase
IACUC: Institutional Animal Care and Use Committee
IAP: The inhibitor of apoptosis proteins
IC50: The half maximal inhibitory concentration
IGF-IR: Insulin like growth factor 1 receptor
IHC: Immunohistochemistry
indels: Insertions and deletions
JH: Johns Hopkins
KEGG: Kyoto Encyclopedia of Genes and Genomes
KIR3DL2: Killer cell immunoglobulin like receptor, three Ig domains and long cytoplasmic tail 2
L: Liter
LFC: Logarithm fold change
LKO: Lentiviral knockout
LOF: Loss of function
Log: logarithm
MAF: Minor allele frequency
MEK: Mitogen-activated protein kinase kinase, MAP2K
mg: Milligram
ml: Milliliter
MN: University of Minnesota
MPNST: Malignant peripheral nerve sheath tumors
MSCV: Murine Stem Cell Virus
NBF: Neutral buffered formalin
NCT: National clinical trials
NF1: Neurofibromatosis Type 1
NF1: Neurofibromin 1
NFRI: NF Research Initiative
ng: nanogram
NIBR: Novartis Institute for Biomedical Research
NIH: National Institutes of Health
NRG: NOD Rag gamma
ns: not significant
NTRK2: Neurotrophic Receptor Tyrosine Kinase 2
Par: Parental
PBS: Phosphate-buffered saline
PCR: Polymerase chain reaction
PD: Pharmacodynamic
PD: Pull down
PDF: Portable document format
PDX: Patient-derived xenograft
PE: Paired-end
pH: Potential of hydrogen
PI: Propidium iodide
PIK3CA: The phosphatidylinositol-4,5-bisphosphate 3-kinase, catalytic subunit alpha
PLK-1: Polo-like kinase 1
pNF: Plexiform neurofibromas
PRC2: Polycomb repressive complex 2
PS: Partially sensitive
PTPN11: Protein tyrosine phosphatase non-receptor type 11
puro: Puromycin
PVDF: Polyvinylidene fluoride
R: Reverse
RABL6A: Rab-like protein 6A
RAF: Rapidly accelerated fibrosarcoma
RAS: Rat sarcoma
RB: Retinoblastoma 1
RBC: Red blood cells
ref: Reference
Res: Resistant
RIPA: Radio-Immunoprecipitation Assay
RNase: Ribonuclease
RNAseq: RNA sequencing
rRNA: Ribosomal ribonucleic acid
RTK: Receptor tyrosine kinases
rtTA: Reverse tetracycline-controlled trans-activator
S: Somatic
S: Sense
SD: Standard deviation
SDS-PAGE: Sodium dodecyl-sulfate polyacrylamide gel electrophoresis
SEM: Standard error of the mean
SHP2: Src homology-2 domain-containing protein tyrosine phosphatase-2
SHP2i: SHP2 inhibitor
shRNA: Short/small hairpin RNA
SKCCC: Sidney Kimmel Comprehensive Cancer Center
SNV: Single nucleotide variant
STR: Short-tandem repeat
SUZ12: Suppressor Of Zeste 12 Protein Homolog
TAZ: Tafazzin
Tet: Tetracycline
TP53: Tumor protein P53
tram: trametinib
TSC2: Tuberous Sclerosis Complex 2
μg: Microgram
μm: Micrometer
μmol: Micromole
v.: versus
VEP: Variant Effect Predictor
VS: Very sensitive
WCL: whole cell lysate
WES: Whole exome sequencing
WNT2B: Wnt family member 2B
WT: Wild-type
WU: University of Washington
YAP: Yes-associated protein

## Supplementary Materials

**Title: CDK4/6 inhibition enhances SHP2 inhibitor efficacy and is dependent upon RB function in malignant peripheral nerve sheath tumors**

**Authors**: Jiawan Wang^1^, Ana Calizo^1^, Lindy Zhang^1^, James C. Pino^2^, Yang Lyu^3^, Kai Pollard^1^, Xiaochun Zhang^3^, Alex T. Larsson^4^, Eric Conniff^5^, Nicolas Llosa^1^, David K. Wood^6^, David A. Largaespada^4^, Susan E. Moody^7^, Sara J. Gosline^2^, Angela C. Hirbe^3^, Christine A. Pratilas^1*^

## Supplementary Figures and Legends

**Fig. S1.**
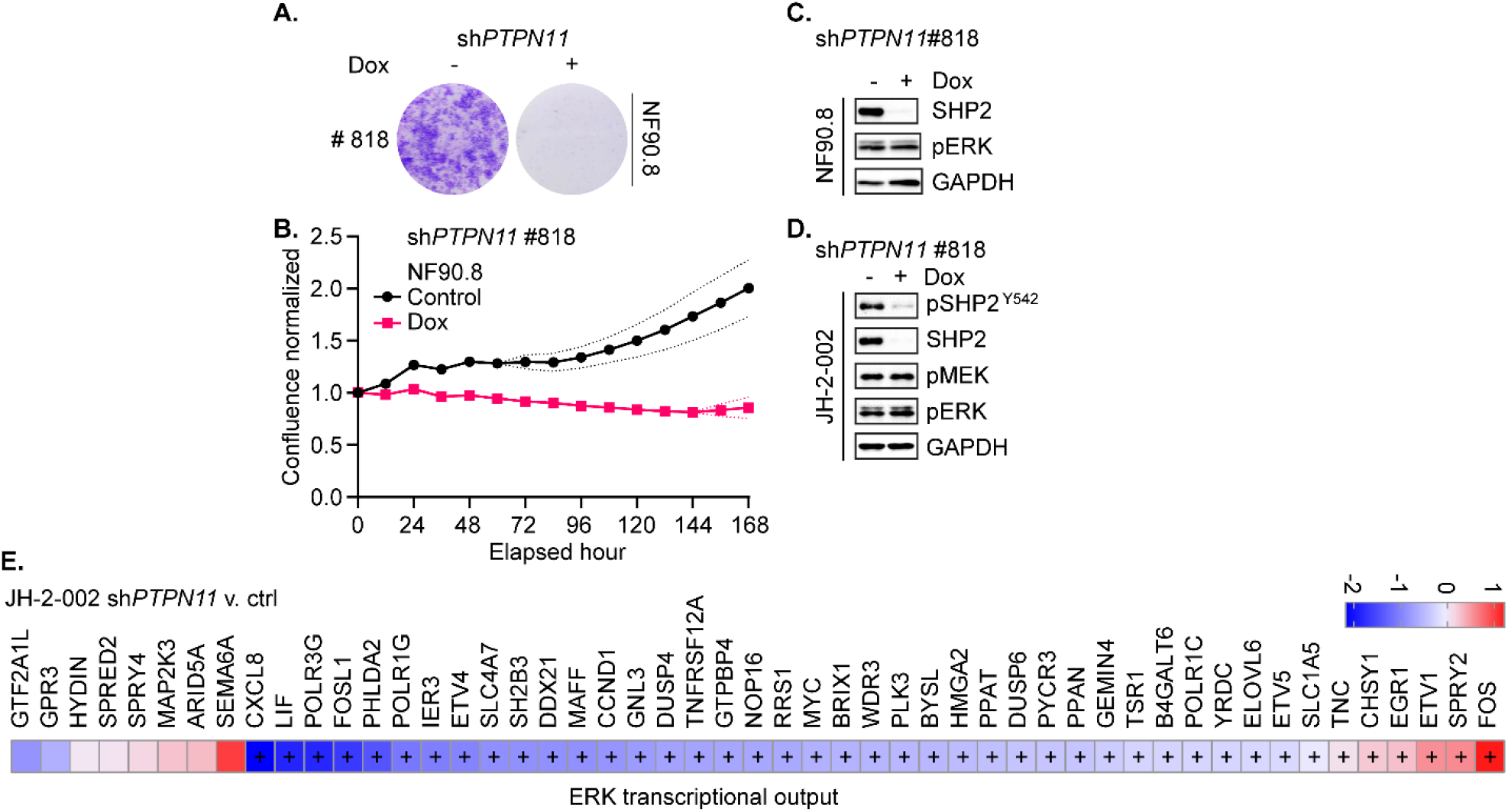
Related to Fig. 1. (**A**) NF90.8 transduced with doxycycline (Dox) inducible constructs expressing sh*PTPN11* #818, was treated with vehicle or 300 ng/ml Dox for two weeks. Cells were fixed with 10% neutral buffered formalin and then stained with crystal violet. (**B)** Cells as in A. were treated with vehicle or 300 ng/ml Dox, and the phase confluence was monitored by Incucyte real-time imaging system, normalized to corresponding 0-hour scan. (**C**) Cells as in A. were treated with vehicle or 300 ng/ml Dox for 72 hours, and the indicated proteins were assessed using immunoblot. (**D**) JH-2-002 as in Fig 1D was evaluated and validated using immunoblot. (**E**) Heatmap related to Fig 1D, demonstrating transcriptional changes of the 51 MEK/ERK signature genes following SHP2/*PTPN11* knockdown. + = *P* adj<0.05.

**Fig. S2.**
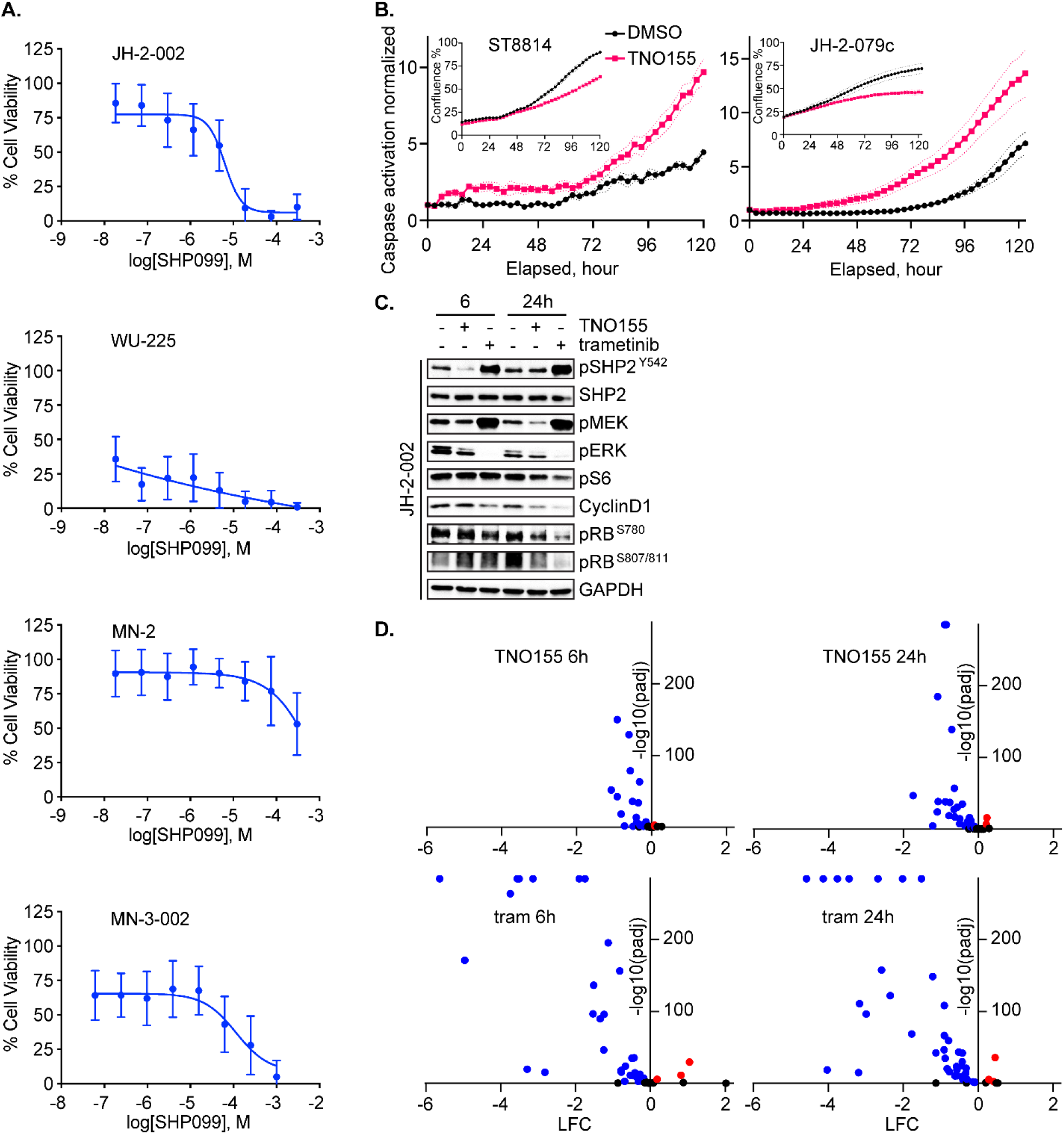
Related to Fig. 2. (**A**) The tumor microtissues from four individual PDX were cultured *in vitro* under 3D conditions and were treated with increasing dose of SHP2i SHP099. (**B**) ST8814 and JH-2-079c were treated with DMSO or 0.3 μmol/L TNO155, together with Incucyte caspase-3/7 dye for apoptosis (Sartorius, #4440) for five days. The normalized caspase activation is shown as the ratio of fluorescence intensity over phase confluence, normalized to corresponding 0-hour scan. (**C**) Corresponding immunoblot to RNAseq samples demonstrating ERK and cell cycle signaling inhibition following 6- and 24-hour treatment with TNO155 and trametinib. (**D**) Volcano plots related to Fig 2I, showing the transcriptional alterations of ERK signaling output comparing TNO155 and trametinib with DMSO control. 53% (27/51) and 63% (32/51) of ERK signature genes after 6- and 24-hour TNO155, and 71% (36/51) and 84% (43/51) of these genes after 6- and 24-hour trametinib, were significantly inhibited (*P* adj<0.05).

**Fig. S3.**
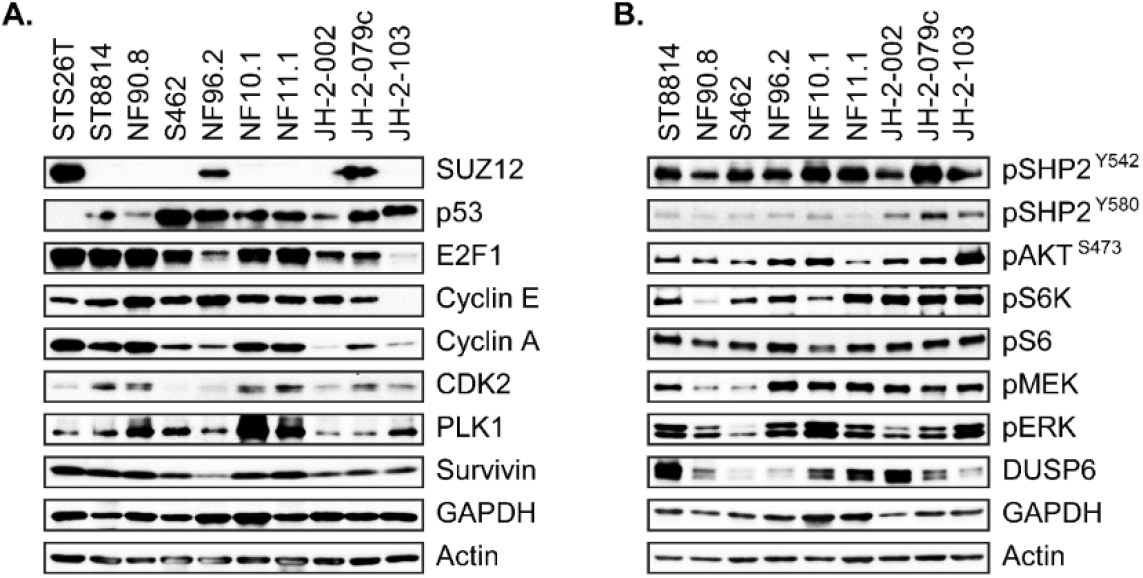
Related to Fig. 3. (**A**) SUZ12 and selected cell cycle regulators were evaluated in the nine NF1-associated and one sporadic (STS26T) MPNST cell lines. (**B**) Baseline level of signaling intermediates in ERK and PI3K/AKT pathways was assessed in the nine NF1-associated MPNST cell lines.

**Fig. S4.**
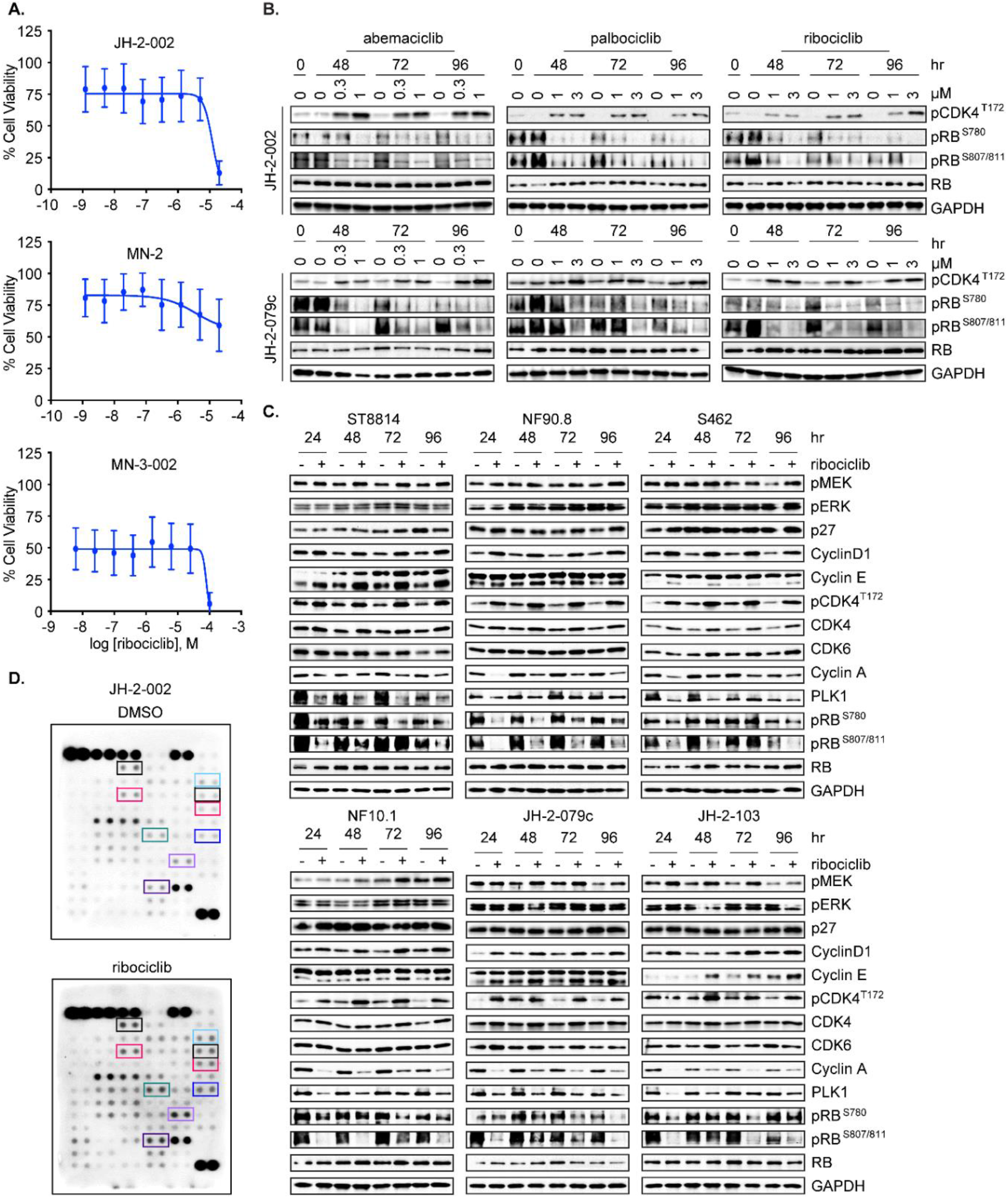
Related to Fig. 4. (**A**) 3D microtissues from three individual PDX were cultured *in vitro* under 3D conditions and were treated with increasing dose of CDK4/6i ribociclib. (**B**) JH-2-002 and JH-2-079c were treated with DMSO, abemaciclib (0.3 and 1 μmol/L), palbociclib (1 and 3 μmol/L), or ribociclib (1 and 3 μmol/L) for 48, 72 and 96 hours. (**C**) Six NF1-MPNST cell lines were treated with DMSO or 1 μmol/L ribociclib over a time course. Signal intermediates in ERK and cell cycle pathways were assessed. (**D**) JH-2-002 cells were treated with DMSO or 1 μmol/L ribociclib for 24 hours. 71 phosphorylated human receptor tyrosine kinases (RTK) were evaluated using human RTK phosphorylation array C1.

**Fig. S5.**
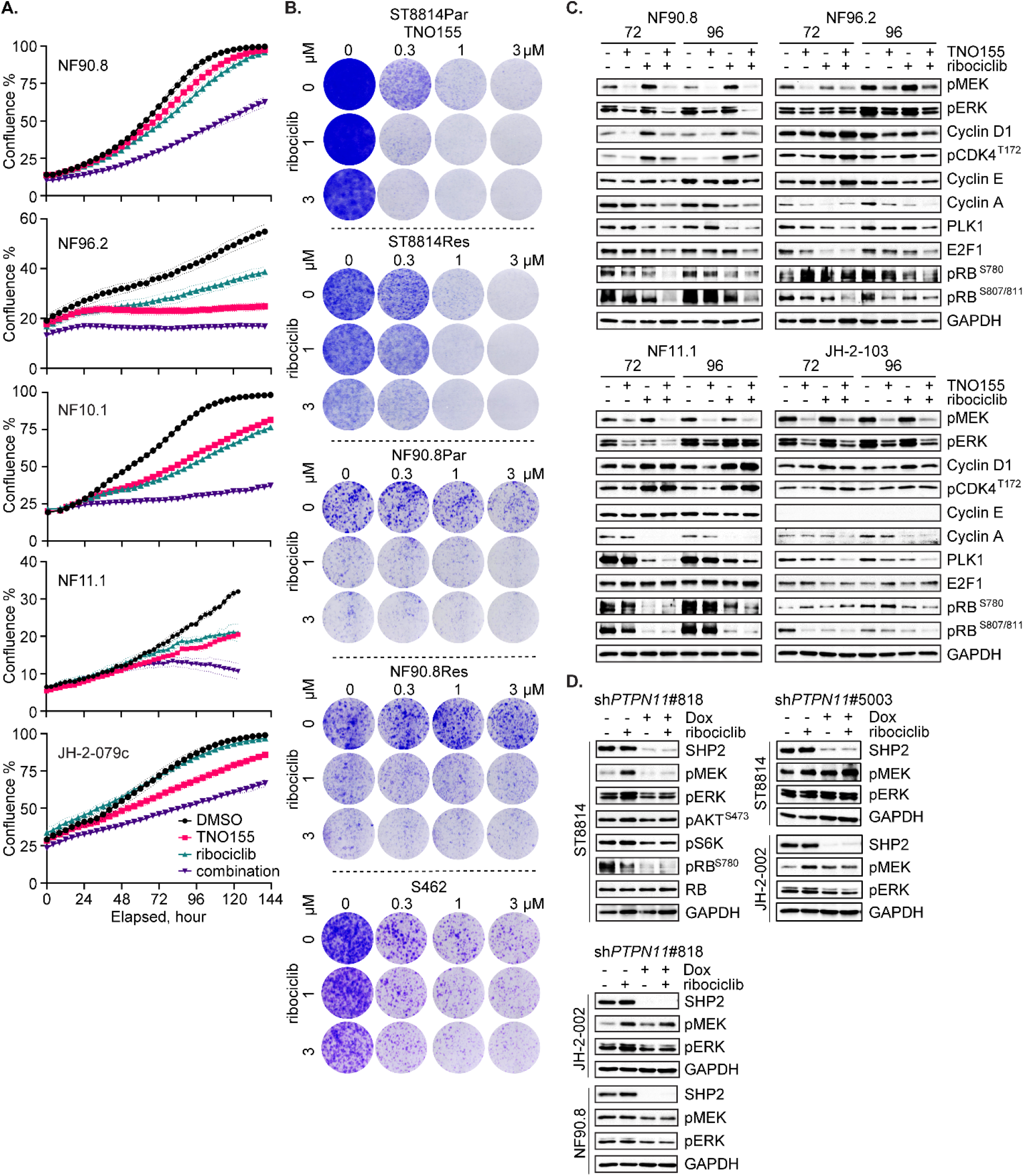
Related to Fig. 5. (**A**) Five NF1-MPNST cell lines were treated with DMSO, TNO155 (0.3 μmol/L), ribociclib (1 μmol/L) and their combination for up to six days. Cell confluence was monitored using Incucyte imaging systems. (**B**) ST8814 and NF90.8 parental and trametinib-resistant cells and S462 were treated with drugs as in Figure 5C for 2-3 weeks, and colony formation was evaluated using crystal violet assay. (**C**) Four NF1-MPNST cell lines were treated with DMSO, 0.3 μmol/L TNO155 and/or 1 μmol/L ribociclib for 72 and 96 hours. ERK signaling and cell cycle regulators were evaluated using immunoblot. (**D**) Three NF1-MPNST cell lines transduced with sh*PTPN11* #5003 or #818 were pretreated with vehicle or 300 ng/ml Dox for 72 hours, followed by treatment with DMSO or 1 μmol/L ribociclib for additional 72 hours. Cell lysates were assessed for expression of the indicated proteins.

**Fig. S6.**
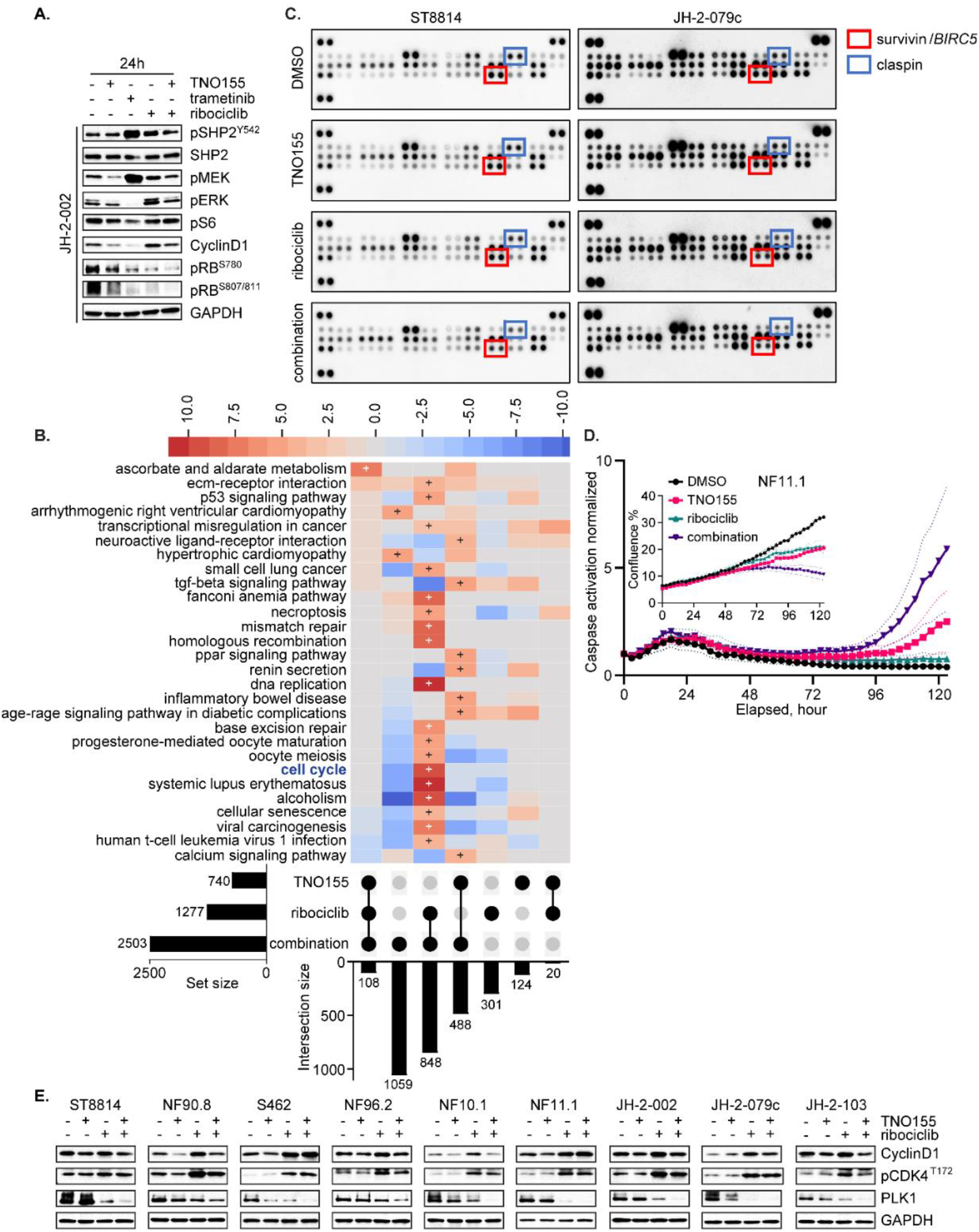
Related to Fig. 6. (**A**) Samples corresponding to RNAseq from JH-2-002 treated with DMSO, 0.3 μmol/L TNO155, 20 nmol/L trametinib, 1 μmol/L ribociclib, and TNO155 plus ribociclib in combination for 24 hours. ERK signaling was assessed using immunoblot. (**B**) Heatmap displaying pathway enrichment against the KEGG database under the 7 intersections on the upset plot in Figure 6A. + = significance. (**C**) ST8814 and JH-2-079c were treated with DMSO, 0.3 μmol/L TNO155 and/or 1 μmol/L ribociclib for 48 hours, and then protein lysates were assessed using human apoptosis antibody array (R&D systems, #ARY009). (**D**) NF11.1 was treated with DMSO, 0.3 μmol/L TNO155 and/ or 1 μmol/L ribociclib, together with Incucyte caspase-3/7 dye for apoptosis (Sartorius, #4440) for five days. The normalized caspase activation is shown as the ratio of fluorescence intensity over phase confluence, normalized to each corresponding 0-hour scan. (**E**) Nine NF1-MPNST cell lines were treated as in C. and the indicated proteins involved in cell cycle signaling were detected using immunoblot.

**Fig. S7.**
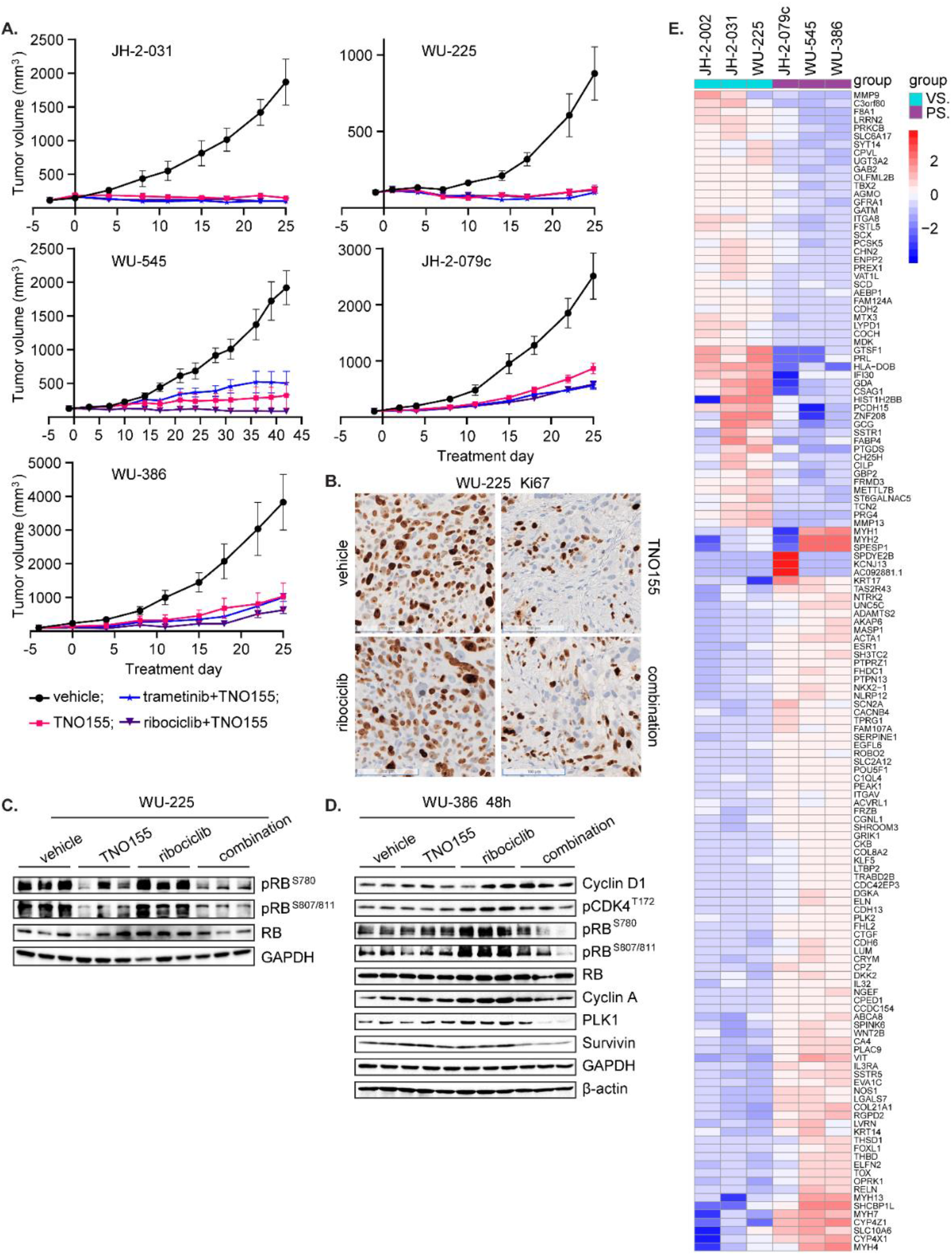
Related to Fig. 7. (**A**) NRG mice bearing five individual PDX were on therapy as in Figure 7A and B, plus one additional arm trametinib (0.15 mg/kg for JH-2-079c and 0.075 mg/kg for others, once daily) in combination with TNO155 (7.5 mg/kg, twice daily). (**B**) NRG mice bearing the PDX WU-225 were on treatment as in Figure 7A, and tumors were collected 4 hours after last dose of 4-week therapy and fixed in 10% NBF. The Ki-67 expression was assessed using IHC. (**C**-**D**) Tumors of each arm from the PDX WU-225 and WU-386 were harvested 4 hours post last dose, 4 weeks on drugs (**C**); or 48 hours post last dose, 3 days on drugs (**D**). The indicated proteins involved in ERK and cell cycle signaling were detected by immunoblot. (**E**) Heatmap displaying 141 differentially expressed genes with adjusted *P* < 0.01 via DeSEQ2 analysis, when comparing very sensitive (VS, better response to TNO155, JH-2-002, JH-2-031 and WU-225) v. partially sensitive (PS, lower response to TNO155, JH-2-079c, WU−545 and WU-386) PDX models.

## Data files S1 to S6 (Excel files)

Table S1_MEK dependent genes_shPTPN11_TNO155_tram

Table S2_DEG_JH-2-002

Table S3_Intersections of samples

Table S4_VS vs PS_de_deseq2

Table S5_Antibody and reagent list

**Table S6.**
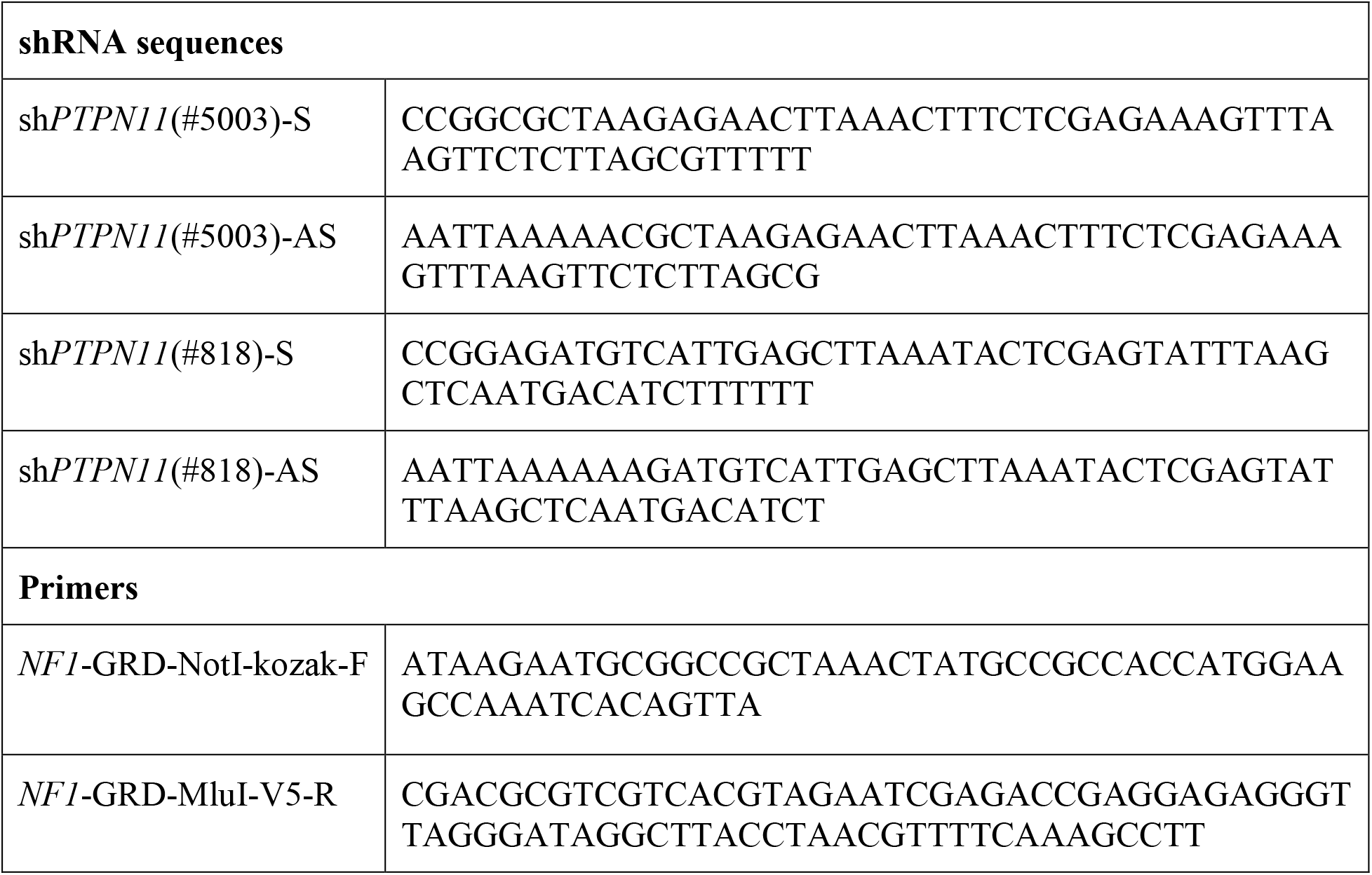
Sequences.

## References and Notes

1. B. S. Ducatman, B. W. Scheithauer, D. G. Piepgras, H. M. Reiman, D. M. Ilstrup, Malignant peripheral nerve sheath tumors. A clinicopathologic study of 120 cases. Cancer 57, 2006–2021 (1986).

2. D. G. Evans et al., Malignant peripheral nerve sheath tumours in neurofibromatosis 1. J Med Genet 39, 311–314 (2002).

3. E. Shurell et al., Gender dimorphism and age of onset in malignant peripheral nerve sheath tumor preclinical models and human patients. BMC Cancer 14, 827 (2014).

4. A. S. Brohl, E. Kahen, S. J. Yoder, J. K. Teer, D. R. Reed, The genomic landscape of malignant peripheral nerve sheath tumors: diverse drivers of Ras pathway activation. Sci Rep 7, 14992 (2017).

5. W. Lee et al., PRC2 is recurrently inactivated through EED or SUZ12 loss in malignant peripheral nerve sheath tumors. Nat Genet 46, 1227–1232 (2014).

6. M. Zhang et al., Somatic mutations of SUZ12 in malignant peripheral nerve sheath tumors. Nat Genet 46, 1170–1172 (2014).

7. T. De Raedt et al., PRC2 loss amplifies Ras-driven transcription and confers sensitivity to BRD4-based therapies. Nature 514, 247–251 (2014).

8. C. Dehner et al., Chromosome 8 gain is associated with high-grade transformation in MPNST. JCI Insight 6, (2021).

9. H. Bhatia et al., Ex vivo to in vivo model of malignant peripheral nerve sheath tumors for precision oncology. (2022).

10. A. Kim, C. A. Pratilas, The promise of signal transduction in genetically driven sarcomas of the nerve. Exp Neurol 299, 317–325 (2018).

11. E. Dombi et al., Activity of Selumetinib in Neurofibromatosis Type 1-Related Plexiform Neurofibromas. N Engl J Med 375, 2550–2560 (2016).

12. A. M. Gross et al. (American Society of Clinical Oncology, 2018).

13. G. B. McCowage et al. (American Society of Clinical Oncology, 2018).

14. R. D. Dodd et al., NF1 deletion generates multiple subtypes of soft-tissue sarcoma that respond to MEK inhibition. Mol Cancer Ther 12, 1906–1917 (2013).

15. W. J. Jessen et al., MEK inhibition exhibits efficacy in human and mouse neurofibromatosis tumors. J Clin Invest 123, 340–347 (2013).

16. E. Jousma et al., Preclinical assessments of the MEK inhibitor PD-0325901 in a mouse model of Neurofibromatosis type 1. Pediatr Blood Cancer 62, 1709–1716 (2015).

17. C. M. Johannessen et al., The NF1 tumor suppressor critically regulates TSC2 and mTOR. Proc Natl Acad Sci U S A 102, 8573–8578 (2005).

18. C. M. Johannessen et al., TORC1 is essential for NF1-associated malignancies. Curr Biol 18, 56–62 (2008).

19. J. Wang, K. Pollard, A. Calizo, C. A. Pratilas, Activation of Receptor Tyrosine Kinases Mediates Acquired Resistance to MEK Inhibition in Malignant Peripheral Nerve Sheath Tumors. Cancer Res 81, 747–762 (2021).

20. J. Wang et al., Combined Inhibition of SHP2 and MEK Is Effective in Models of NF1-Deficient Malignant Peripheral Nerve Sheath Tumors. Cancer Res 80, 5367–5379 (2020).

21. Y. N. Chen et al., Allosteric inhibition of SHP2 phosphatase inhibits cancers driven by receptor tyrosine kinases. Nature 535, 148–152 (2016).

22. C. Fedele et al., SHP2 Inhibition Prevents Adaptive Resistance to MEK Inhibitors in Multiple Cancer Models. Cancer Discov 8, 1237–1249 (2018).

23. R. J. Nichols et al., RAS nucleotide cycling underlies the SHP2 phosphatase dependence of mutant BRAF-, NF1- and RAS-driven cancers. Nat Cell Biol 20, 1064–1073 (2018).

24. A. Prahallad et al., PTPN11 Is a Central Node in Intrinsic and Acquired Resistance to Targeted Cancer Drugs. Cell Rep 12, 1978–1985 (2015).

25. S. Bunda et al., Inhibition of SHP2-mediated dephosphorylation of Ras suppresses oncogenesis. Nature communications 6, 8859 (2015).

26. J. Bendell et al., Intermittent dosing of RMC-4630, a potent, selective inhibitor of SHP2, combined with the MEK inhibitor cobimetinib, in a phase 1b/2 clinical trial for advanced solid tumors with activating mutations of RAS signaling. European Journal of Cancer 138, S8–S9 (2020).

27. J. L. Kohlmeyer et al., RABL6A Is an Essential Driver of MPNSTs that Negatively Regulates the RB1 Pathway and Sensitizes Tumor Cells to CDK4/6 Inhibitors. Clin Cancer Res 26, 2997–3011 (2020).

28. J. Hagen et al., RABL6A promotes G1-S phase progression and pancreatic neuroendocrine tumor cell proliferation in an Rb1-dependent manner. Cancer Res 74, 6661–6670 (2014).

29. M. Bentires-Alj et al., Activating mutations of the noonan syndrome-associated SHP2/PTPN11 gene in human solid tumors and adult acute myelogenous leukemia. Cancer Res 64, 8816–8820 (2004).

30. K. Pollard et al., A clinically and genomically annotated nerve sheath tumor biospecimen repository. Sci Data 7, 184 (2020).

31. C. A. Pratilas et al., (V600E)BRAF is associated with disabled feedback inhibition of RAF-MEK signaling and elevated transcriptional output of the pathway. Proc Natl Acad Sci U S A 106, 4519–4524 (2009).

32. J. Cai et al., High-risk neuroblastoma with NF1 loss of function is targetable using SHP2 inhibition. Cell Rep 40, 111095 (2022).

33. F. Xing et al., Concurrent loss of the PTEN and RB1 tumor suppressors attenuates RAF dependence in melanomas harboring (V600E)BRAF. Oncogene 31, 446–457 (2012).

34. C. Giacinti, A. Giordano, RB and cell cycle progression. Oncogene 25, 5220–5227 (2006).

35. K. Munger, D. L. Jones, Human papillomavirus carcinogenesis: an identity crisis in the retinoblastoma tumor suppressor pathway. J Virol 89, 4708–4711 (2015).

36. G. W. Demers, E. Espling, J. B. Harry, B. G. Etscheid, D. A. Galloway, Abrogation of growth arrest signals by human papillomavirus type 16 E7 is mediated by sequences required for transformation. J Virol 70, 6862–6869 (1996).

37. S. I. Gharbi et al., Crystal structure of active CDK4-cyclin D and mechanistic basis for abemaciclib efficacy. NPJ Breast Cancer 8, 126 (2022).

38. L. R. Pack, L. H. Daigh, M. Chung, T. Meyer, Clinical CDK4/6 inhibitors induce selective and immediate dissociation of p21 from cyclin D-CDK4 to inhibit CDK2. Nat Commun 12, 3356 (2021).

39. S. Paternot, B. Colleoni, X. Bisteau, P. P. Roger, The CDK4/CDK6 inhibitor PD0332991 paradoxically stabilizes activated cyclin D3-CDK4/6 complexes. Cell Cycle 13, 2879–2888 (2014).

40. D. B. Solit et al., BRAF mutation predicts sensitivity to MEK inhibition. Nature 439, 358–362 (2006).

41. M. Alvarez-Fernandez, M. Malumbres, Mechanisms of Sensitivity and Resistance to CDK4/6 Inhibition. Cancer Cell 37, 514–529 (2020).

42. X. Chen et al., Discovery of a Novel Src Homology-2 Domain Containing Protein Tyrosine Phosphatase-2 (SHP2) and Cyclin-Dependent Kinase 4 (CDK4) Dual Inhibitor for the Treatment of Triple-Negative Breast Cancer. J Med Chem 65, 6729–6747 (2022).

43. C. Liu et al., Combinations with Allosteric SHP2 Inhibitor TNO155 to Block Receptor Tyrosine Kinase Signaling. Clin Cancer Res 27, 342–354 (2021).

44. G. J. Lee et al., Maximizing the therapeutic potential of SHP2 inhibition with rational combination strategies in tumors driven by aberrant RAS-MAPK signaling. 79, 1322–1322 (2019).

45. N. T. Shifrin et al., Dual inhibition of SHP2 and CDK4/6 leads to immunological memory and immune-mediated anti-tumor activity in a mouse syngeneic model of breast cancer. 80, 2837–2837 (2020).

46. G. A. Martin et al., The GAP-related domain of the neurofibromatosis type 1 gene product interacts with ras p21. Cell 63, 843–849 (1990).

47. A. M. Bolger, M. Lohse, B. Usadel, Trimmomatic: a flexible trimmer for Illumina sequence data. Bioinformatics 30, 2114–2120 (2014).

48. H. J. a. p. a. Li, Aligning sequence reads, clone sequences and assembly contigs with BWA-MEM. (2013).

49. D. C. Koboldt et al., VarScan 2: somatic mutation and copy number alteration discovery in cancer by exome sequencing. Genome Res 22, 568–576 (2012).

50. S. Kim et al., Strelka2: fast and accurate calling of germline and somatic variants. Nat Methods 15, 591–594 (2018).

51. D. Benjamin et al., Calling somatic SNVs and indels with Mutect2. 861054 (2019).

52. K. Ye, M. H. Schulz, Q. Long, R. Apweiler, Z. Ning, Pindel: a pattern growth approach to detect break points of large deletions and medium sized insertions from paired-end short reads. Bioinformatics 25, 2865–2871 (2009).

53. W. McLaren et al., The Ensembl Variant Effect Predictor. Genome Biol 17, 122 (2016).

54. R. Patro, G. Duggal, M. I. Love, R. A. Irizarry, C. Kingsford, Salmon provides fast and bias-aware quantification of transcript expression. Nat Methods 14, 417–419 (2017).

55. M. I. Love, W. Huber, S. Anders, Moderated estimation of fold change and dispersion for RNA-seq data with DESeq2. Genome Biol 15, 550 (2014).

56. W. Hadley, Ggplot2: Elegrant graphics for data analysis. (Springer, 2016).

57. K. A. Cummins, A. L. Crampton, D. K. Wood, A High-Throughput Workflow to Study Remodeling of Extracellular Matrix-Based Microtissues. Tissue Eng Part C Methods 25, 25–36 (2019).

58. A. L. Crampton, K. A. Cummins, D. K. Wood, A high-throughput microtissue platform to probe endothelial function in vitro. Integr Biol (Camb) 10, 555–565 (2018).

59. C. Stringer, T. Wang, M. Michaelos, M. Pachitariu, Cellpose: a generalist algorithm for cellular segmentation. Nat Methods 18, 100–106 (2021).

